# Assessment of Menstrual Health Status and Evolution through Mobile Apps for Fertility Awareness

**DOI:** 10.1101/385054

**Authors:** Laura Symul, Katarzyna Wac, Paula Hillard, Marcel Salathé

## Abstract

For most women of reproductive age, assessing menstrual health and fertility typically involves regular visits to a gynecologist or another clinician. While these evaluations provide critical information on an individual’s reproductive health status, they typically rely on memory-based self-reports, and the results are rarely, if ever, assessed at the population level. In recent years, mobile apps for menstrual tracking have become very popular, allowing us to evaluate the reliability and tracking frequency of millions of self-observations, thereby providing an unparalleled view, both in detail and scale, on menstrual health and its evolution for large populations. In particular, we were interested in exploring the tracking behavior of the app users and their overall observation patterns in an effort to understand if they were consistent with previous small-scale medical studies. We also investigated whether their precision allowed the detection and estimation of ovulation timing, which is critical for reproductive and menstrual health. Retrospective self-observation data were acquired from two mobile apps dedicated to the application of the sympto-thermal fertility awareness method, resulting in a dataset of more than 30 million days of observations from over 2.7 million cycles. The analysis of the data showed that up to 40% of the cycles in which users were seeking pregnancy had recordings every single day. With a modeling approach using Hidden Markov Models to describe the collected data and estimate ovulation timing, it was found that follicular phases average duration and range were larger than previously reported, with only 24% of ovulations occurring at days 14 to 15, while the luteal phase duration and range were in line with previous reports, although short luteal phases (10 days or less) were more frequently observed (in up to 20% of cycles). The digital epidemiology approach presented here can help to lead to a better understanding of menstrual health and its connection to women’s health overall, which has historically been severely understudied.

## Introduction

A broad diversity of fertility awareness methods (FAMs) has been developed in the past century^1,2^, primarily designed to help couples manage fertility and family planning. Modern methods developed in the last quarter of the 20^th^ century take advantage of the precise description of menstrual variation of the basal body temperature (BBT) or waking temperature, taken with a thermometer with a 0.01C or 0.5F precision, cervical mucus quality and quantity, vaginal sensation, and cervical position^3–6^. These methods have defined a set of rules that allows the identification of the fertile window around ovulation, so that couples can adapt their sexual behavior according to their reproductive objectives^7–9^. The sympto-thermal method, which combines BBT and cervical mucus observations, is arguably amongst the most reliable FAM for family planning^1,2,4,10^. Recently, a number of mobile apps have been developed by private organizations to facilitate FAM tracking. Some of these apps provide their users with automatized interpretation with regard to the opening and closing of the fertility window^11^. Over the past few years, an increasing number of women, estimated at over 200 million in 2016^12^, have started using these apps, contributing to the accumulation of menstrual-related data (Fig 1) from a diverse population of users at different stage of life (Fig. 2A, Table 1, Methods).

**Table 1:**
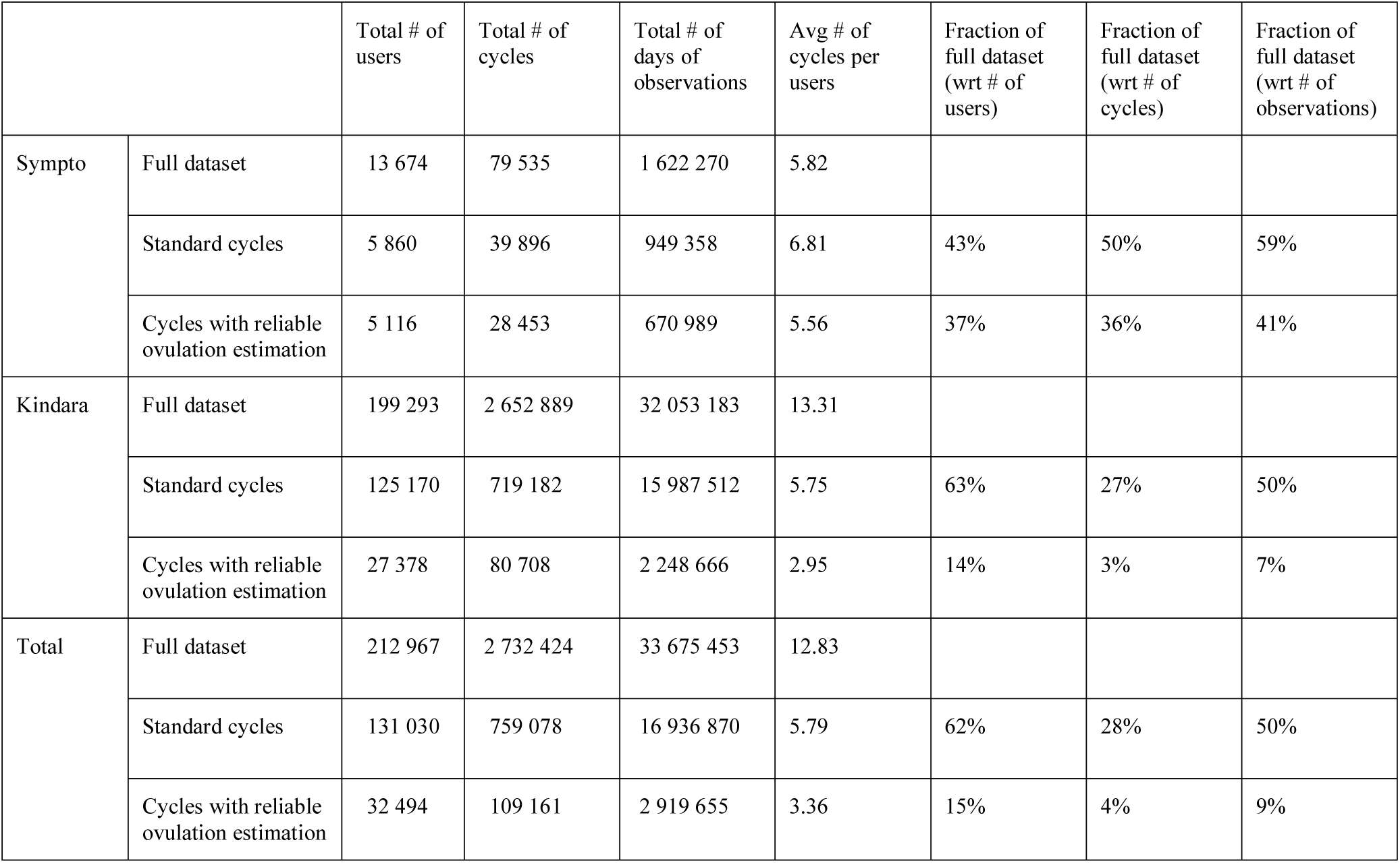
number of observations, cycles and users. Number of users, cycles and days of observations. In a single day, a user can log up to 7 observations, *i.e.* one in each of the tracking categories available to users, see Table 2.

|  |  | Total # of users | Total # of cycles | Total # of days of observations | Avg # of cycles per users | Fraction of full dataset (wrt # of users) | Fraction of full dataset (wrt # of cycles) | Fraction of full dataset (wrt # of observations) |
| --- | --- | --- | --- | --- | --- | --- | --- | --- |
| Sympto | Full dataset | 13 674 | 79 535 | 1 622 270 | 5.82 |  |  |  |
|  | Standard cycles | 5 860 | 39 896 | 949 358 | 6.81 | 43% | 50% | 59% |
|  | Cycles with reliable ovulation estimation | 5 116 | 28 453 | 670 989 | 5.56 | 37% | 36% | 41% |
| Kindara | Full dataset | 199 293 | 2 652 889 | 32 053 183 | 13.31 |  |  |  |
|  | Standard cycles | 125 170 | 719 182 | 15 987 512 | 5.75 | 63% | 27% | 50% |
|  | Cycles with reliable ovulation estimation | 27 378 | 80 708 | 2 248 666 | 2.95 | 14% | 3% | 7% |
| Total | Full dataset | 212 967 | 2 732 424 | 33 675 453 | 12.83 |  |  |  |
|  | Standard cycles | 131 030 | 759 078 | 16 936 870 | 5.79 | 62% | 28% | 50% |
|  | Cycles with reliable ovulation estimation | 32 494 | 109 161 | 2 919 655 | 3.36 | 15% | 4% | 9% |

**Figure 1:**
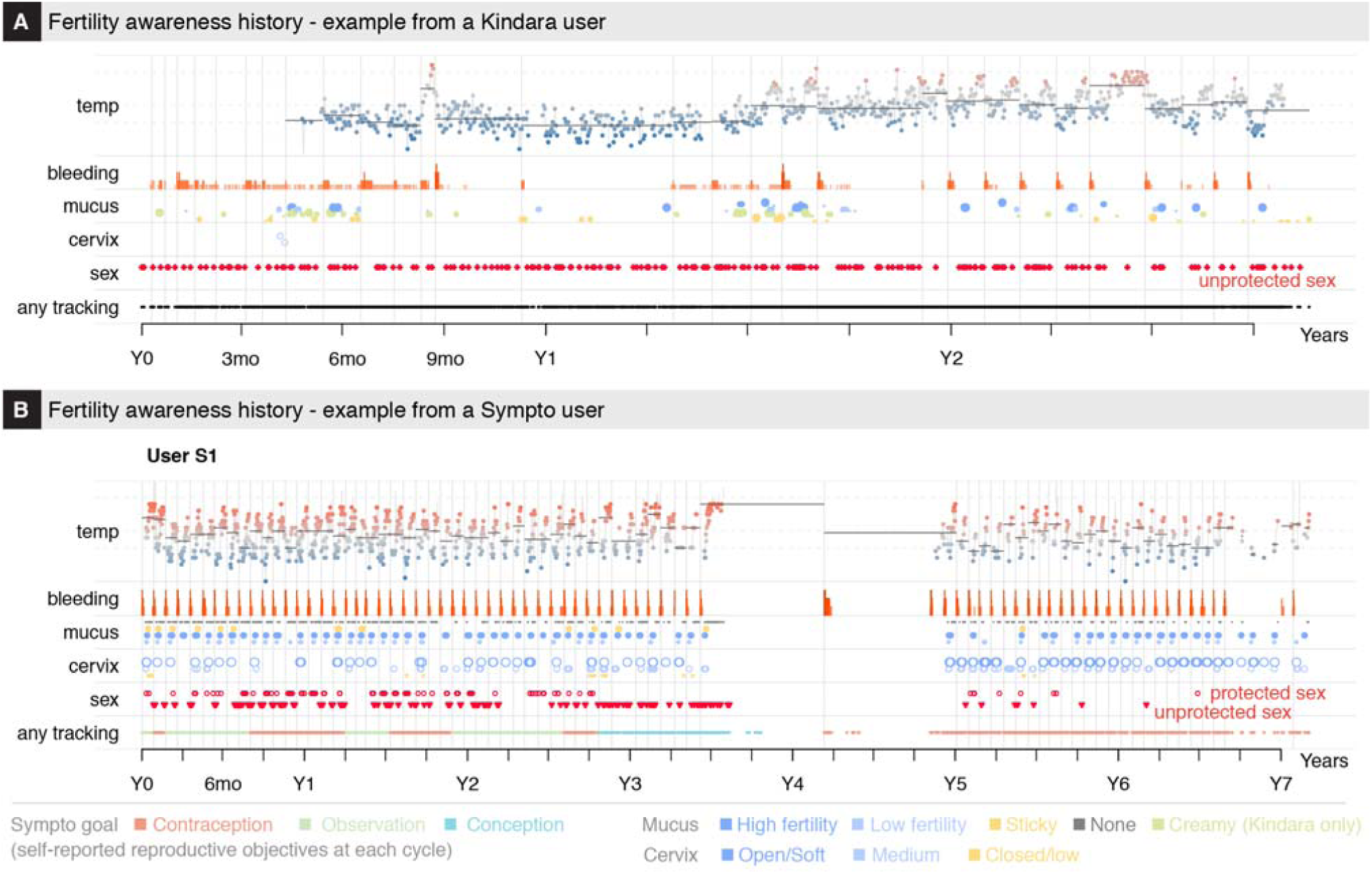
Menstrual history of two app users. Menstrual history of two long term Kindara (**A**) and Sympto (**B**) users. Time is shown in years as relative to the first observation of each user. Kindara user is seeking to achieve pregnancy and show a long anovulatory episode during which her overall temperature profile is lower. She returns to more regular, ovulatory cycles in her last year of tracking, as indicated by the bleeding frequency and the temperature profiles. The Sympto user has used the app to avoid pregnancy and observe her cycle for almost 3 years, before trying to conceive, which she likely achieves after 9 cycles (her reported reproductive objective switches from “contraception” to “conception” – line “any tracking” at the bottom). 9 months later, the user reports bleeding, which likely indicates post-partum bleeding (lochia). After another 9 months, probably as she stops breastfeeding, she logs menstrual observations and returns to using the app to avoid pregnancy.

**Figure 2:**
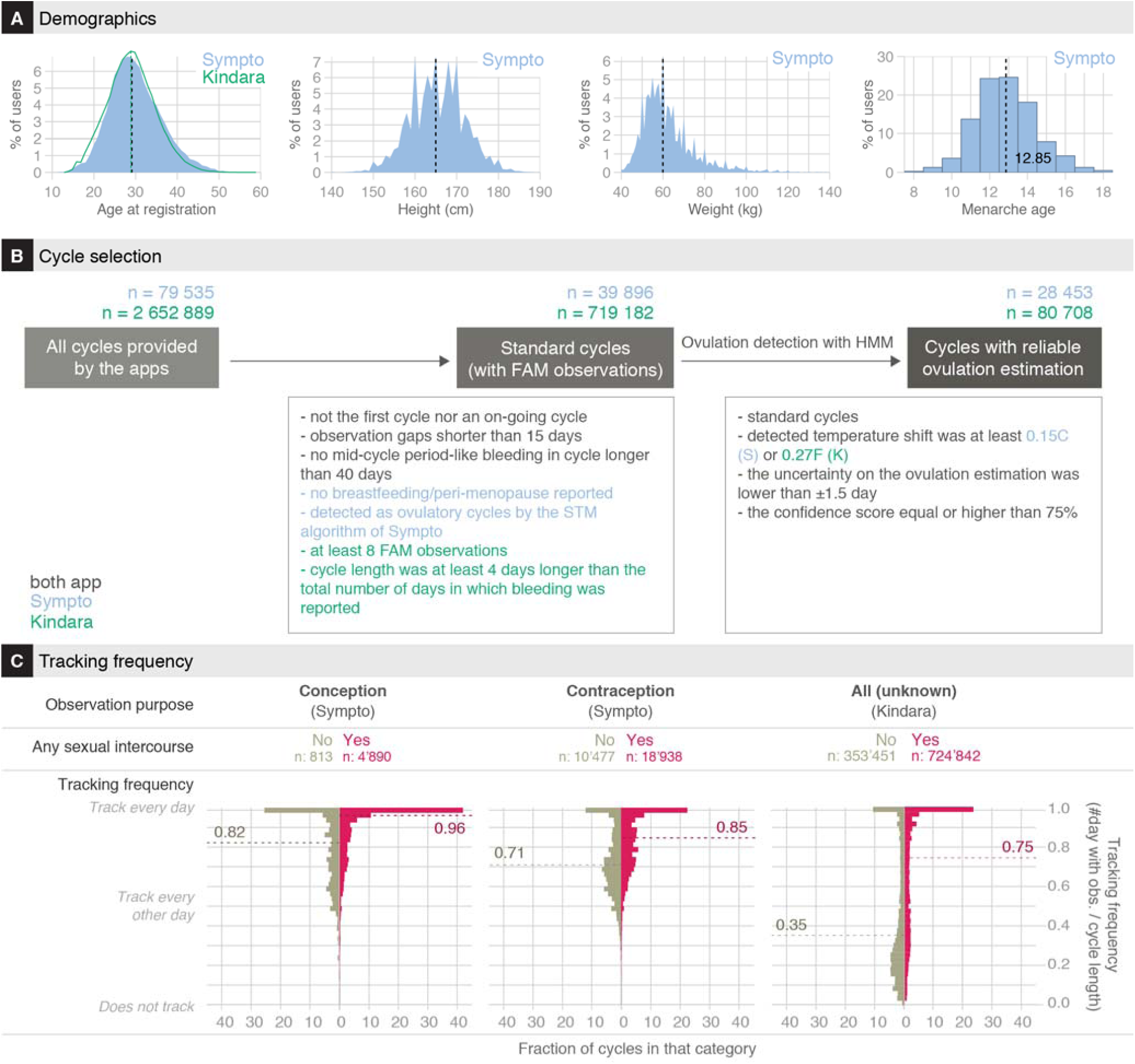
Demographics and tracking behavior of users. **(A)** Users’ age at registration (left), reported height (middle-left), weight (middle-right) and menarche age of users (right). **(B)** Cycle selection flowchart. Methods provide extensive description of the inclusion/exclusion criteria. *Standard cycles* are finished, complete cycles, typical of a non-pregnant, non-peri-menopausal, non-nursing user, that have at least 8 days with FAM observations (Kindara) or that are detected as ovulatory cycles according to the Sympto implementation of the STM rules. *Cycles with reliable ovulation estimation* are cycles for which the ovulation day could be reliably estimated by the HMM framework developed for this study (Methods). **(C)** Cycle-specific tracking frequencies (top: Sympto, bottom: Kindara). 39,896 (Sympto) + 719,182 (Kindara) *standard cycles* were used (Methods). Dashed lines indicate median values.

A few studies have evaluated some of these apps in terms of user experience or the accuracy of the scientific information provided to their users^13,14^ or regarding their ability to accurately indicate the opening and closing of the fertile window^11,15^. Other studies^16–18^ have evaluated the contraceptive efficacy of the app Natural Cycles; this app based on a proprietary algorithm only takes body temperature into account^16–18^ and these studies were authored by one of the app founders and did not provide a description of the tracked data. In the last two years, only a few studies have used datasets from women’s health applications, such as Clue, to describe the association between pre-menstrual symptoms and sexually transmitted infections^19^ or to develop machine learning methods suited to study rhythmic human behavior^20^ or predict pregnancy^21^.

However, fertility awareness body signs, as tracked easily via affordable mobile applications, have not yet been extensively described or studied and it is unclear how app users are reporting these signs, as well as whether the reported observations are consistent with the conclusions of previous smaller-scale medical studies^6,22,23^. Moreover, there are no statistical framework to detect ovulation from these self-tracked data, which would be useful to leverage the potential of these data to study fertility, accurately predict pregnancy chances and to overall evaluate the potential impact of fluctuating hormones on the course of chronic diseases^24^.

Here, we were interested in evaluating the potential of the data collected via apps for the assessment of menstrual health, both at the individual level and at the population level, and, in the long run to help enable better clinical-decision-making processes. Two retrospective datasets that were described by the app providers as representative of their active users population (Methods), were acquired from the apps Kindara (**K**) and Sympto (**S**)^11^ (Fig. S1A,B, Table 1). Both apps offer free and paid versions of their app, but all data used in this study can be tracked on their free version. Privacy policies of the two apps explicitly state that users’ data might be transferred to academic institutions for research purposes with the motivation to support studies that could potentially accelerate future development of fertility awareness methods. Both apps offer similar FAM tracking options but differ in their design and user experience (Fig. S1AB, Table 2). Kindara is primarily marketed to women who wish to achieve pregnancy and does not provide feedback to users in terms of the opening or closing of their fertile window. Sympto is marketed as a family planning tool that can be utilized to plan or avoid a pregnancy. The Sympto app provides feedback to their users based on their observations, indicating when they are potentially fertile, very fertile or infertile. The key differences between these two apps are (i) the automatic- (**S**) *vs* user- (**K**) interpretation of observations, (ii) the per-cycle (S) *vs* per-user (K) definition of fertility goals users wish to achieve, (iii) the criteria for the onset of a new cycle, i.e. fresh bleeding after ovulation (**S**) *vs* self-assessed or automatic (K), and (iv) the precision at which users can report their observations (Suppl. Mat.).

**Table 2:** Reported observations. Tracking options available to users of the Sympto and Kindara app. Kindara offers more granularity and categories for reporting mucus, cervix and vaginal sensation. Provided that they primarily market users who wish to achieve pregnancy, they also offer the option to track insemination. Sympto considers withdrawal as unprotected sex and does not offer that option to their user.

| type | Sympto |  | Kindara |  |
| --- | --- | --- | --- | --- |
|  | unit/categories | max precision / subcategories | unit/categories | max precision / subcategories |
| BBT | Celsius | 0.05 | Fahrenheit | 0.01 |
| BBT time | daytime | 1/2h | daytime | minute |
| questionable temp | (not recorded) |  | logical |  |
| mucus | NA |  | NA |  |
|  | no mucus |  | no mucus |  |
|  | little amounts of creamy mucus or not very stretchable mucus |  | creamy | little/medium/lots |
|  | large amounts of egg-white like, watery, very stretchable mucus |  | egg-white like | little/medium/lots |
|  |  |  | watery | little/medium/lots |
|  | sticky mucus |  | sticky | little/medium/lots |
| cervix | NA |  | height | low/medium/high |
|  | closed, firm, low |  | firmness | firm/medium/soft |
|  | medium |  | openness | closed/medium/open |
|  | open, soft, high |  |  |  |
| vaginal sensation | NA |  | NA |  |
|  | dry |  | dry |  |
|  | wet |  | dry sticky |  |
|  | very wet |  | wet moist |  |
|  |  |  | wet lubricate |  |
| sex | protected |  | protected |  |
|  | unprotected |  | unprotected |  |
|  |  |  | withdrawal |  |
|  |  |  | insemination |  |

## Results

### Users demographics: the typical FAM app user is 30, has a healthy BMI, and lives in a European or North American country

The two apps target different populations. Most Kindara users are based in the US and are trying to achieve pregnancy, while Sympto users mainly reside in Europe and use the app primarily to avoid pregnancy. Users of these two apps are found in over 150 countries, covering 5 continents, but the vast majority of them are located in Europe and in the Americas. User ages span the reproductive life of women, from the onset of their sexual activity to menopause, with an overrepresentation of users in their late 20s and early 30s (Fig. 2A, left, Table 3). For some users, additional information is available, including their birth year, and, for Sympto users only, their reported weight, height and age at menarche (Fig 2A, Table 3).

**Table 3:** users demographics. Users demographic information.

|  | **Mean ± standard deviation** | **Fraction of users with available information (%)** |
| --- | --- | --- |
| **Age** | 29.9 ± 6.4 (Sympto)<br>29.2 ± 5.9 (Kindara) | 81 (Sympto)<br>13 (Kindara) |
| **Age at Menarche** | 12.9 ± 1.6 (Sympto) | 79 (Sympto) |
| **Height (in cm)** | 161 ± 21(Sympto) | 80 (Sympto) |
| **Weight (in kg)** | 62 ± 13 (Sympto) | 80 (Sympto) |
| **BMI** | 23 ± 5 (Sympto) | 80 (Sympto) |

The height and weight distribution of Sympto users (Fig. 2A, top and bottom right, data not available for Kindara users, Table 3) shows median values of 60 kg and 165 cm. Both distributions present peaks at round values such as 160 or 165 cm indicating that users often report approximate values (for example, 160 cm rather than 159 or 161 cm). This has often been observed in previous studies using self-reported values and these mild inaccuracies of self-reported values have usually been found to only slightly affect the overall distributions ^25^. The median BMI of Sympto users is around 20, which is considered healthy for women (Fig S1C, Table 3). Information such as users’ level of education, marital or social status, parity or particular health conditions are unknown.

### Regular users log their observations at a high frequency

The tracking behavior of regular FAM users during their usual cycles, which here are referred to as “*standard cycles*” (Fig 2B, Methods) is highly variable and depends on the family planning objectives of the users (Fig 2C). For an idealized ∼28-day cycle, FAM-relevant body signs need to be recorded for at least 8-12 days of each cycle to detect the changes related to ovulation, *i.e.* at a tracking frequency of at least ∼ 43%. However, most users using the apps for their FAM tracking report their observations for over 16 days per cycle. In cycles where users choose to record sexual intercourse (65% (S) – 75% (K) of *standard cycles*), tracking frequency is increased, with over 40% of cycles being tracked every single day when seeking pregnancy (Fig. 2C, S1D), sometimes for several months or years in a row (Fig. 1).

Tracking frequencies varied between the two apps (Fig 2C), partly in relationship to the design of the apps; Kindara doesn’t provide user interpretation of the fertility window allowing for sporadic tracking, whereas missing data in Sympto precludes an accurate fertility assessment, potentially leading the user to discontinue use of the app if they are unable or unwilling to track consistently.

### Reported fertility awareness body signs show clear patterns at the user population level

Confident that users regularly logged observations (Fig. 2C) during *standard cycles*, we sought to characterize general patterns in the observations and frequency of the different FAM body signs and investigate whether they were consistent with previous studies^5,6,9,26,27^. As cycle durations vary by several days, as illustrated in Fig. 3A, and given that the duration of the luteal phase (after ovulation) has been shown to vary less than the follicular phase (before ovulation)^28,29^, ovulation-related observations (BBT, mucus, cervix, vaginal sensation) are shown from the end of each cycle (Fig. 3B-D, S2). A clear shift of about 0.36°C/0.7°F in BBT between the mid-follicular phase and the mid-luteal phase is observed (Fig 3B, S2A), consistent with previous observations on a cohort of much smaller size^26^. BBT showed a decrease at the end of the cycle, as light bleeding or spotting was reported (Fig. 3BC).

**Figure 3:**
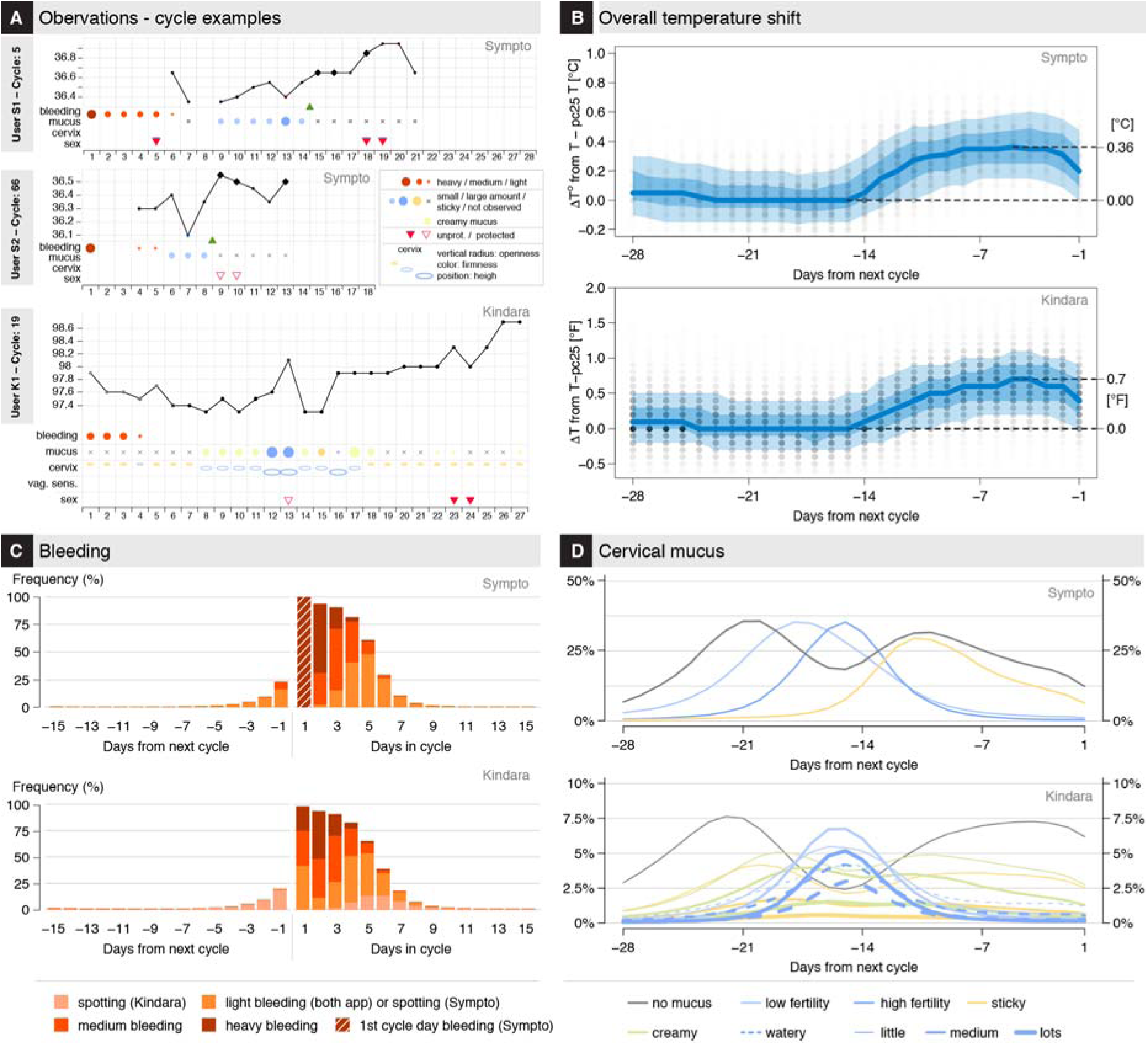
User observations overview. **(A)** Examples of observations: the 5^th^ tracked cycle (top) and 66^th^ cycle (middle) cycle of two different Sympto users. Observations of the 19^th^ cycle (bottom) of a Kindara user. **(B)** ΔBBT (variation from the 25% percentile of temperature in this cycle) values are shown on each day of the cycle, from the end of the cycle. Opacity of the dots reflects the number of observations. The median value: thick blue line. 10, 25, 75 and 90 percentiles of ΔBBT: translucent blue bands. **(C)** Frequency of bleeding observations, for the end (left) and beginning (right) of cycles. The Sympto app only starts a new cycle on the first recording of heavy bleeding (score 3/3, dark red) after a post-ovulatory infertile phase, thus all cycles present heavy bleeding at the start of the cycle (hashed dark red bar). **(D)** Frequency of cervical mucus observations from the end of cycles (top: S, bottom: K). (Kindara) Little quantity of watery mucus (dashed line) and little or medium quantity of egg-white like mucus (solid line) are considered as ‘low fertility’ mucus (light blue) while large quantities of egg-white like and medium or large quantities of watery mucus are considered as ‘high fertility’ mucus (dark blue) (B-D) 39,896 (S) + 719,182 (K) *standard cycles* were used (Methods)

In an ovulatory cycle, it is well established that cervical mucus is produced in higher quantity and with a higher stretchiness in the days leading up to ovulation^5,6,9,27^, which seems to be observed by users tracking their cervical mucus (85-90% (S) and 40-45% (K) of cycles) (Fig. 3D).

### Estimation of ovulation day reveals the diversity of menstrual timing

Previous studies have shown that the combination of BBT and cervical mucus variations were reliable, although not perfect, proxies for the detection of ovulation^8,23,27,30^. We therefore decided to define a mathematical framework (HMM) to derive an estimate of the most likely day of ovulation with reliability indicators to reflect the uncertainty of conflicting or unexpected observation patterns (Fig. 4A, S3,4,6, Methods). Missing temperature records have been found to alter the precision of the ovulation estimation to a slightly greater extent than missing cervical mucus reports (Fig S6D, Suppl. Mat.).

**Figure 4:**
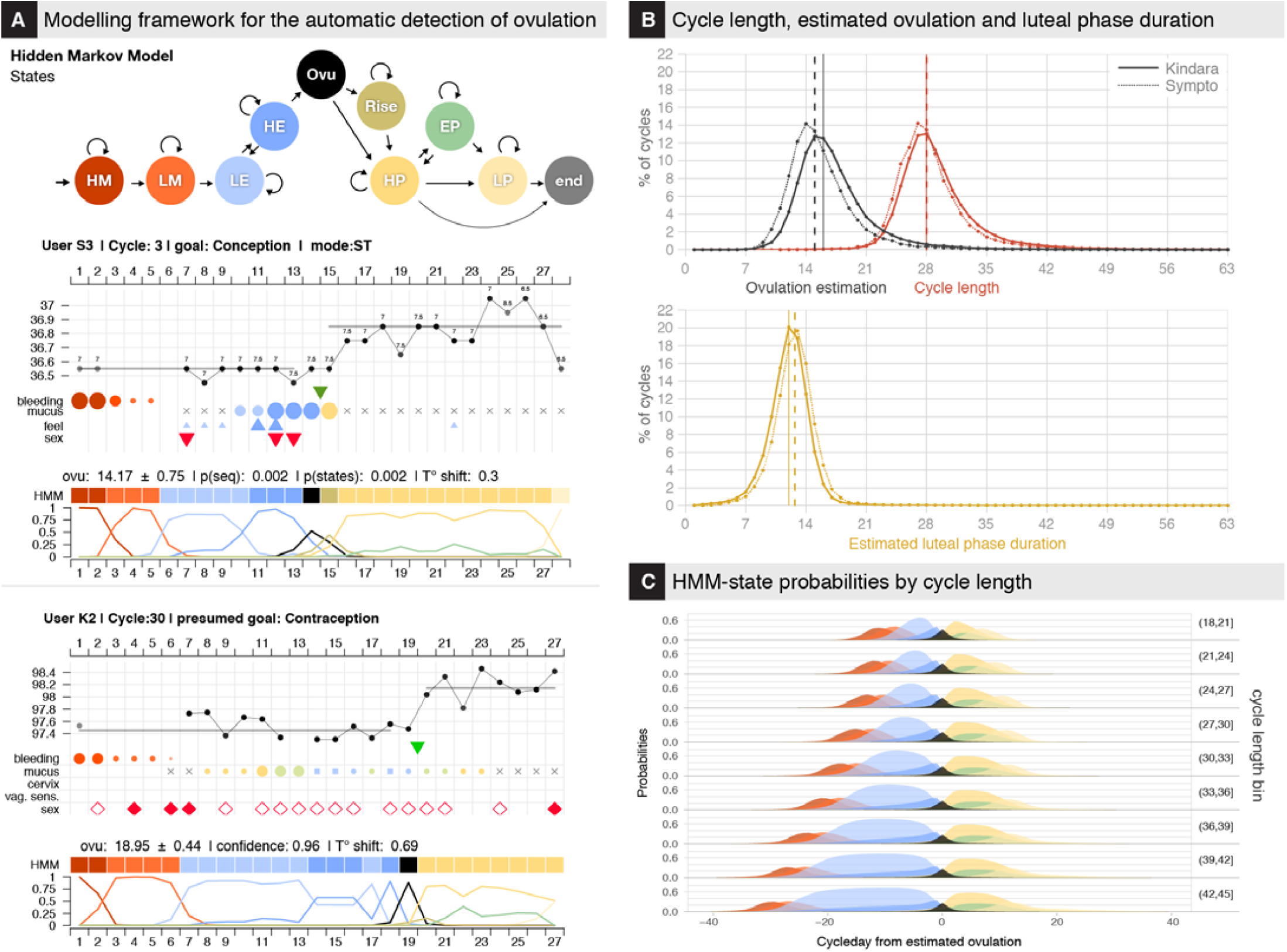
Modeling framework for the estimation of ovulation and menstrual states. **(A)** Modeling framework for the estimation of ovulation timing. (Top) Schematics of the 10-states HMM which discretizes the menstrual hormonal events (HM: Heavy Menses, LM: Light Menses, LE: Low Estrogen, HE: high Estrogen, Ovu: Ovulation, Rise: progesterone/BTT Rise, HP: High Progesterone, EP: Estrogen Peak in luteal phase, LP: Low Progesterone). Arrows indicate possible state-transition; arrow thickness is not representative of actual transition probabilities (Methods). (Bottom) Examples of menstrual state estimation for the 2rd and 3rd cycle of 2 users. (Top of each chart) Original user observations as in Fig. 2A. (Middle of each chart) Colored squares HMM-labeled line) represent the most likely sequence of HMM states given the observations (Methods). (Bottom of each chart) Normalized probabilities of each state on each day of the cycle (Methods). **(B)** (Top) Cycle length and estimated ovulation day. (Bottom) Luteal phase duration, computed as the number of days between the ovulation day (excluded) and the1st day of the next cycle (excluded). Vertical lines indicate median values. 80,708 (K) + 24,119 (S) *cycles with reliable ovulation estimation* were used (Methods). **(C)** Average estimated state probabilities by cycle-day counting from estimated ovulation aggregated by total cycle length (in bins of 3 units) for all *cycles with reliable ovulation estimation.*

These estimations allowed the comparison, for *cycles with reliable ovulation estimation* (109,161 cycles, Methods), of the cycle length distribution to those of estimated day of ovulation and of the duration of the luteal phase (*i.e.* post-ovulation) (Fig. 4B). Cycle length distribution is asymmetrical around the typical 27 to 28 days, with a heavy tail on longer cycles. Similarly, the distribution of the follicular (*i.e.* prior to ovulation) phase duration (or ovulation time) is asymmetrical as well, with a median value of 16 days, and 90% of ovulations occurring between day 10 and day 24. Only ∼24% of ovulations occurred on days 14 to 15 of the cycle.

Luteal phase duration distribution, which is also asymmetrical, presents however a skew for smaller values and a smaller standard deviation (Fig. 4BC, S4BC). Median values were 12 (K) and 13 (S) days, which is in line with a previous study that used fertility monitors^31^ but shorter than values reported in studies that used luteinizing hormone (LH) peak for timing of ovulation^29,32^. About 35% of cycles have a luteal phase duration of 12 to 13 days, while ∼20% of cycles had a luteal phase duration smaller than or equal to 10 days, which represents a higher proportion than reported in a previous epidemiological study^29^.

Overall, the comparison with previous studies of the cycle phases duration and range shows that the follicular phase and the whole cycle length have higher mean values and larger ranges than what was previously observed, while the luteal phase duration and range was closer to those found in previous studies^28,29,31,33,34^ (Fig. S5). The larger observed mean and range of the follicular phase and the cycle length can partially be explained by the differences in inclusion/exclusion criteria – for example, some previous studies excluded long cycles (Table S9) – and by the ovulation estimation methods, but also probably by the fact that this study uses cycles from a much larger population and is thus able to capture a higher diversity of menstrual patterns.

Interestingly, the cycle phases distributions were slightly different when considering the data from the two apps. These differences might be due to biases found in the user population, especially for users seeking pregnancy that could be at higher risk of sub-fertility if assumed that they start tracking after they have already tried to get pregnant for several months (Fig S4C); however, these data on user behaviors around fertility seeking are not available for Kindara users.

## Discussion

This study’s goal was to describe and explore the suitability of datasets collected through two mobile applications (Kindara and Sympto) supporting Fertility Awareness Method (FAM) tracking for the assessment of menstrual health in general, both at the individual level and at the population level. We found that the average tracking frequency of users that utilize the apps FAM tracking, *vs* basic period tracking, is higher than the minimum required to detect changes associated with ovulation. In particular, if users rely on the app for their family planning, *i.e.* if they log sexual intercourses (protected or unprotected), the tracking frequency is increased, with up to 40% of cycles having recordings every single day when the user’s objective is to achieve pregnancy. The reported observations (BBT, cervical mucus changes, etc.) are overall aligned with expected patterns of FAM-related body signs, showing these apps enable hundreds of thousands of users across Europe and North America to follow their fertility and ovulation patterns. However individual cycles often present noisier profiles and missing data are a frequent concern. To partly alleviate these issues, the mathematical framework (HMM) used in this study discretizes the menstrual cycle in independent successive biologically-relevant states and allows the estimation of ovulation timing along with uncertainty indicators. A large variation in the ovulation time and in the luteal phase duration was found, with larger ranges than previously described in other studies^29,31,33,36^ that relied on much smaller populations but that used biomarkers which might offer a greater precision for the estimation of ovulation time.

The strength of this study lies in the scale and precision of the datasets, as a variety of fertility patterns are captured, and as users track the evolution of their cycles at a high frequency over long intervals of time. It also provides a non-proprietary and replicable mathematical method to infer biological states, and in particular to estimate the timing of ovulation, from fertility awareness self-tracked data. The most obvious potential limitation of this study comes from the origin of these retrospective data: a self-selected possibly biased population, limited medical and general information on users, irregular observation patterns and little control on assessing the validity of the observations, in particular with regard to cervical mucus tracking. While the tracking frequency limitation can be alleviated through strict selection of users and cycles (Methods), all other limiting factors might have introduced biases in the present analysis. Prospective studies on selected cohorts with appropriate follow-up and information provided to users will provide higher quality data, which could then be used for comparison.

Based on the current findings, it appears that self-tracking of FAM-related body signs provides an affordable means to evaluate the status and evolution of menstrual health, given that these observations require only a precise thermometer, and that providers of these and of other apps offer free simplified versions. These long term and yet very precise recordings support the idea that the menstrual cycle, like other biological rhythms, is a vital sign whose variations inform about overall health status^37,38^. The digital epidemiology approach^39^, where patients collect data themselves through digital means, can in this context represent a powerful method to investigate menstrual health and its connection to women’s health at the population level^34^ in a field that has historically been severely understudied^40^.

We foresee that future studies will use self-tracked data to quantify infertility or daily pregnancy chances based on reported FAM body signs and user’s history. Models could also be established to investigate potential sub-fertility causes (anovulation, recurrent early pregnancy losses, etc.) based on the fertility signs and user’s sexual behavior. More generally, such data and tracking apps, combined with tracking of other coexisting symptoms, enable the exploration of the menstrual dimension of the course of chronic diseases^24,41^. Such studies would highly benefit from additional, sometimes already existing, tracking options in the apps such as pregnancy validation (for example reports of pregnancy tests results) or a prompt to the user to label a tracking pause such that it can reliably be differentiated from a pregnancy.

It is likely that users of such applications already have an increased awareness of their cycles, and this study suggests that these digitally self-tracked observations potentially present an opportunity to facilitate the dialog between patients and their clinicians, helping them to make informed decisions based on quantified indicators. The current and future development of evidence-based digital tools for menstrual health monitoring could positively impact women’s health.

## Methods

Extended Materials and Methods can be found in the Supplementary Materials.

To briefly summarize the methodology used in this study: datasets were first filtered to keep cycles of users using the apps for fertility awareness purposes, i.e. identify their fertility window. Data were then summarized to describe the overall observation patterns. Finally, a Hidden Markov Model (HMM) was defined and used to detect ovulation time and assess the reliability of this estimation.

### Mobile phone applications and data acquisition

Two de-identified retrospective datasets were acquired from the Symptotherm foundation (www.sympto.org; Switzerland) and Kindara (www.kindara.com; US) upon receiving ethical approval from the Canton Geneva ethical commission (CCER Genève, Switzerland), study number 2017-02108. These two apps were selected as they both ranked high in a study comparing the performances of apps marketed to avoid pregnancy using FAMs^11^, as their privacy policies specified the use of their deidentified datasets for research purposes and as their user pools were very large or diverse geographically and culturally. Sympto has been released in 2008 and is available worldwide in 8 languages (English, French, German, Italian, Spanish, Polish, Russian and Bulgarian). Kindara has been released in 2012 and is available worldwide in English. Both apps de-identified their datasets before transferring them to the authors. Both apps are available on iOS and Android platforms and are available as free or paid apps. All features used in this study are available in the free versions of the apps. Kindara provided a random subset of their overall pool of users with at least 4 logged cycles (199 293 users, 2 652 889 cycles) while Sympto provided observations from their long-term users (at least 4 cycles tracked with the app) and from users who provided their weight, height and menarche age (13 674 users, 79 535 cycles). A description of the datasets fields is provided in Table 2.

### Selection criteria for users and cycles

Given that these are self-tracked data, missing data is a frequent issue, and many cycles within the datasets provided by the app were not suitable for the analyses of this study. First cycles were filtered to remove any unfinished or uncomplete cycles or cycles in which fertility awareness body signs were not observed. Kept cycles are labelled as “standard cycles” (see flowchart, Fig 2B). Then, the HMM was used to estimate ovulation and, for the rest of the analysis, only cycles in which ovulation could reliably be estimated were kept (Fig 2B). Below are the inclusion/exclusion criteria for these cycle categories.

#### Standard cycles

(Sympto: 39,896 cycles; Kindara: 719,182 cycles) denote cycles of regular users of the apps in which FAM body signs have been logged. Typically, cycles with long tracking gaps or in which only the period flow was logged were excluded.

- (S&K) not the first cycle of a user nor an on-going cycle

- (S&K) observation gaps were no longer than 15 days within a given cycle

- (S&K) at least one FAM body sign (BBT or cervical mucus or cervix position) was recorded

- (S&K) no mid-cycle period-like bleeding was detected when the cycle was longer than 40 days

- (S) defined as ovulatory cycles by the STM algorithm of Sympto, *i.e.* in which the fertile window could be closed.

- (K) at least 8 FAM observations were reported

- (S) no breastfeeding was reported or peri-menopause was declared

- (K) cycle length was at least 4 days longer than the total number of days in which bleeding was reported

#### Cycles with reliable ovulation estimation

(Sympto: 28,453 cycles; Kindara: 80,708)

Criteria summary:

*- standard cycles*

- detected temperature shift was at least 0.15C (S) or 0.27F (K) (Suppl. Mat.).

- the uncertainty on the ovulation estimation was lower than ±1.5 day (Suppl. Mat.).

- the confidence score, which is related to acceptable amount of missing data in the ovulatory period, was equal or higher than 75% (Suppl. Mat.).

### Observations decoding and ovulation timing estimation with HMM

The FAM body-signs are considered to reflect the hormonal changes orchestrating the menstrual cycles. The study was focused on understanding the extent to which these tracked cycles were consistent with previously described menstrual cycle physiologic changes, and the extent to which it was thus possible for app users to estimate timing of ovulation. Hidden Markov Models (HMM) are one of the most suitable mathematical frameworks to estimate ovulation timing, due to their ability to uncover, from observations, latent phenomenon, which in this use include the cascade of hormonal events across the menstrual cycle. HMM have also been previously used for analysis of menstrual periodicity^20^. A 10-states HMM, in which each state is a particular phase of the menstrual cycle (Fig. 4A top, S3A, Suppl. Mat.), was defined, and with decoding algorithms (Viterbi – Backward-Forward) was used to estimate the ovulation time, the uncertainty on this estimation, and a confidence score that accounts for missing observation and variation in temperature taking times.

A set of stringent criteria were established, and included: the uncertainty of the ovulation estimation (≤ ±1.5 days); the magnitude of the temperature shift (≥ 0.15 C); and the confidence score of the observations (≥ 0.75) to discriminate between cycles for which the estimations could be trusted (*cycles with reliable ovulation estimation*) and those where the observations did not allow for a reliable estimation of the ovulation day (Fig S4A, Suppl. Mat.). These strict criteria lead to the exclusion of ∼40% (Sympto) and ∼ 89% (Kindara) of the *standard cycles* that were initially selected. In total, 28,453 (Sympto) + 80,708 (Kindara) *cycles with reliable ovulation estimation* have been used for the subsequent analyses (Suppl. Mat.).

### Model description

The Hidden Markov Model (HMM) as implemented in this study describes a discretization in 10 states of the successive hormonal events throughout an ovulatory menstrual cycle. The HMM definition includes the probabilities of observing the different FAM reported body signs in each state (emission probabilities) and the probabilities of switching from one state to another (transition probabilities). Emission probabilities were chosen to reflect observations previously made in studies that tested for ovulation with LH tests or ultrasounds^6,8,27^, while transition probabilities were chosen in a quasi-uniform manner (Suppl. Mat.). The ovulation estimations were robust to changes in transition probabilities but not to variations in emission probabilities (Fig S6, SI), indicating that this simple framework is suitable to detect ovulations in cycles of any length, and potentially including pregnancies, relying primarily on users’ observations.

Once the model was defined, the Viterbi and the Backward-Forward algorithms^42^ were used to calculate the most probable state sequence for each cycle (Suppl. Mat.) and thus to estimate ovulation timing, i.e. the most likely day of the cycle in which the HMM is in the state “ovulation”. An uncertainty of the estimation has also been computed as the standard deviation of the distribution of probabilities for the state ‘ovulation’, which can be interpreted as the confidence interval in days for the time of ovulation estimation (Suppl. Mat.). Finally, a confidence score was defined to account for missing observations and variation in temperature taking time in a window of ∼5 days around the estimated ovulation day (Suppl. Mat.).

### HMM states

The 10 states, defined as a discretization of the hormonal evolution across the cycle (further details in Suppl. Mat.), are:

HM: onset of the menses and the heavy/medium flow of fresh blood;

LM: days of light bleeding or spotting that conclude menstruations;

LE: Low Estrogen;

HE: High Estrogen; Ovu: Ovulation;

Rise: Temperature rise associated with rise in progesterone production;

HP: High Progesterone;

EP: Estrogen Peak in luteal phase;

LP: Low Progesterone;

*End: Artificial state for the end of each cycle.*

## Acknowledgments

The authors are deeply grateful to all Kindara and Sympto users whose data have been used for this study and to the Symptotherm foundation and Kindara company. In particular, we thank Dr. H. Wettstein, C. Bourgeois, V. Salonna, T. Newcomer, T. Baras, C. Allémann, P. Ducoeurjoly and F. Goddyn for sharing their experience, references and for fruitful discussions. We thank S. Holmes, C. Droin and G. Lazzari for discussion on the mathematical modeling.

## Author contributions

L.S. initiated and conceived the study, analyzed the data and designed the figures, L.S., K.W., P.H. and M.S. wrote the manuscript. All authors discussed the results and implications and commented on the manuscript at all stages.

## Data and code availability

While the privacy policies and terms of usage of the two apps allow the sharing of their de-identified users’ data with third parties for research purposes, they do not allow public sharing of the raw datasets. Aggregated values necessary for the production of the figures as well as the code of the analyses are available at https://lasy.github.io/FAM-Public-Repo/.

