## Supplementary materials for "Assessment of Menstrual Health Status and Evolution through Mobile Apps for Fertility Awareness"

**This PDF file includes:**

Supplementary Text and extended Materials and Methods

Figs. S1 to S5 and Supplementary Figure Legends

Tables S1 to S10
References for SI

Privacy policies of the two applications

**Other supplementary materials for this manuscript include the following:**

Aggregated datasets for each figure and supplementary figures panels.

***Supplementary Text and extended Materials and Methods***

#### Data acquisition

In order to get access to datasets, we contacted the Symptotherm foundation (www.sympto.org) and Kindara (www.kindara.com). The Symptothem foundation was founded in 2001 in Switzerland by H. Wettstein and C. Bourgeois and its mission is to facilitate the diffusion and access to the STM. In 2008, they released the mobile and web app “Sympto”. Kindara was founded in 2011 in the US (Boulder, Colorado, USA) and released their app “Kindara” in 2012 with the mission to support women in achieving their fertility goals. They both ranked in top positions by the only published study to have compared the performance of apps marketed to avoid pregnancy (1). Sympto belongs to the category of apps which provides an automatic detection of the fertile window and Kindara to the category of apps which lets the users interpret their own observations (1).

Upon reaching an agreement with these two organizations and receiving ethical approval from the Canton Geneva ethical commission (CCER Genève, Switzerland), study number 2017-02108, observations of their users were de-identified by the two organizations and subsequently shared with our team for the analyses presented in this study.

Kindara provided a representative sample of their overall pool of users with at least 4 logged cycles while Sympto provided observations from their long-term users and users who provided their weight, height and menarche age.

**App design main differences**

(i) Kindara does not provide any automatic interpretation in terms of potential fertility while the Sympto app relies on an algorithm that implements the rules of the Sympto-thermal method (7, 9), which combines BBT and cervical mucus observations, for providing users with indication regarding the opening and closing of their fertility window.

(ii) Kindara users are invited to define their fertility goal (“Get pregnant”, “avoid pregnancy” or “track my period”) when they start using the app. They can later change this option, but their goal is not recorded as a cycle-specific feature. The Sympto app, however, prompt their users at the beginning of each cycle, to set their goal for that cycle which can be one of the following four: “I wish to become pregnant” (Conception), “I wish to avoid pregnancy” (Contraception), “I wish to observe my cycle” (Observation), “I am open to whatever comes” (Whatever comes). According to their goal, the app displays different icons for the fertile window and motivational messages are adapted.

(iii) Kindara provide the option to let their users decide when a new cycle starts while Sympto will automatically start a new cycle on the first day of “heavy/fresh bleeding” (bleeding score 3/3) but only if an ovulation has been detected by their algorithm in the previous cycle. This has implication in differentiating between potentially anovulatory cycles (or cycles with missing or noisy observations) and ovulatory cycles. However, Sympto’s decision to only start a new cycle with a bleeding score of 3/3 creates an artefact on the bleeding distribution on the first day of each cycle. Also, Kindara let differentiate light bleeding (score 1/3) from spotting, which results in slightly different frequency distributions of light/spotting bleeding.

(iv) Finally, the last main difference in their tracking options is found in the precision at which users can report the quality and quantity of their cervical mucus and of their cervix position. Sympto has defined 3 mucus categories while Kindara has defined 4 categories, which can each vary in 3 intensities. Regarding the cervix tracking options, Kindara let their users track its openness (closed, medium, open), firmness (firm, medium, soft) and height (low, medium, high) as three independent parameters while Sympto has summarized cervix configuration in 3 categories (“closed, firm, low”, “medium” and “open, soft, high”). See Table 2 for details about the reported observations from both apps.

We observed that the app specificities have a crucial impact on the tracking frequency distribution (Fig 2C). Indeed, the algorithm embedded in the Sympto app does not trigger a new cycle if ovulation has not been detected, and about 10-12 observations are necessary on average for a typical ~28-day cycle (tracking frequency ~ 40%). If an ovulation has not been detected, the Sympto app consider that cycle as an anovulatory episode. The tracking frequency of regular Kindara users present a bigger pool at lower tracking frequencies, especially when users do not log sexual intercourse and thus do not rely on the app for important life choices. We did not split cycles tracking with Kindara per goal, as this information was not available per cycles.

**Collected data**

**Bleeding**

Bleeding is reported by app users as heavy (score 3/3), medium (2/3) or light (1/3). Kindara has an additional “spotting” category. When investigating the bleeding patterns reported by apps users (Fig. 3C), we found that Kindara users report their most abundant bleeding on the 2nd day of their cycle. For users of both apps, heavy bleeding frequency rapidly decreases as frequency of medium (score 2/3, bright red) and then of light bleeding (score 1/3, orange) increases. Light bleeding and/or spotting is increasingly observed at the end of cycles, as the next menses are about to start. On the day before the onset of their menses, up to 25% of users report light bleeding or spotting. Spotting, mid-cycle or ovulation-related bleeding is rarely reported. A small fraction (10-15%) of users do not track their bleeding patterns beyond the 1st or 2nd day. As mentioned above, all cycles in Sympto must start with “heavy” bleeding (Fig. 3C).

**Basal Body Temperature (BBT)**

Basal body temperature is the temperature taken at wake with a thermometer that is precise to 0.01C or 0.05 F. Users of both apps are encouraged to report the time at which they took their temperature to potentially detect outliers or aberrant values.

In addition, Sympto users are prompted to report which orifice they use for temperature measurements for a given cycle, which allowed us to explore the temperature distribution by orifice (Fig S2A). We found that the overall temperature distribution is ~ 0.15C degree lower when taken in the mouth than in the vagina or anus. BBT also showed more distinct distributions between pre- and post-ovulation when taken in the anus.

Thus, to evaluate the overall BBT shift across the menstrual cycle, we subtracted the lowest 25% percentile temperature of each cycle from the temperature profile. This allowed us to minimize the inter-individual variability and to account for the measurement noise and for the chosen orifice (mouth, vagina or anus) for temperature taking (Fig. S2A).

**Cervical mucus**

Cervical mucus is collected by FAM users either by inserting one or two clean fingers in their vagina up to their cervix, where it is produced, or by examination of what is found at the vulva (potential small delay).

Counselors of the Symptotherm foundation have shared in private conversations that many users felt unsure of their mucus observations, especially during the first months of using the method.

Here, we found that users start observing their cervical mucus as their period ends, with mucus quality being reported by 85-90% of Sympto users and 40-45% of Kindara users in their late follicular phase and early luteal phase. Users frequently use the option of reporting ‘no mucus observed’ instead of leaving that body sign unreported. After the menses, we first observe a wave of ‘no mucus’ (gray), closely followed by a wave of creamy (beige in the figure) or egg-white like mucus in small quantity (light blue in the figure), then larger quantities of egg-white like or watery mucus (blue, solid (egg-white) and dashed (watery) lines). After presumed ovulation, opaque sticky mucus (yellow in the figure) is observed as well as another wave of ‘no mucus’, both decreasing until the next menses. Creamy mucus or small amounts of sticky mucus (Fig. 2D, bottom) is observed throughout the menstrual cycle, potentially indicating confusion between vaginal desquamation and cervical mucus itself or perhaps indicating signs of mild vaginitis or cervicitis/STD.

**Cervix attributes**

Users of both apps are also given the option to report the status of their cervix. In particular, Kindara users are invited to report the height of their cervix in the vaginal cavity, the openness of the cervix and its softness. The cervix has been reported by Kindara users to be more open and softer in the days leading to ovulation (Fig. S2B). Similarly, although the signal is less strong, cervix is reported to be higher before ovulation (Fig. S2B). Sympto has combined these three attributes into a 3-state scale going from closed/firm/low to open/soft/high and show a similar modification of the cervix attributes round the days preceding ovulation (Fig. S2B). About 10% of Sympto cycles and 5% of Kindara cycles have cervix attributes reported.

**Vaginal Sensation**

The vaginal sensation reflects how users perceive their inner lubrication, moistness.

In the collected data, we found that only few users (35% of Sympto cycles, less than 0.05% of Kindara users - this might reflect that the option has been added later) report their vaginal sensation and rather rely on their cervical mucus to assess their potential fertility status. Users of both apps however report lubricated sensation in the days leading up to ovulation (Fig. S2C).

#### Selection criteria for users and cycles

***Standard cycles*** (Sympto: 39,896 cycles; Kindara: 719,182 cycles)

Standard cycles denote usual cycles of regular users of the apps. We excluded cycles that were unfinished or in which there were large observation gaps (which could indicate a tracking pause or a pregnancy). We also excluded the first and last (unfinished) cycles of each user and cycles for which only the bleeding was reported. Furthermore, we also checked for artificial ‘double cycles’ among those tracked with Sympto. Indeed, as described above, Sympto only triggers a new cycle if its algorithm first had closed the fertility window (i.e. in brief: if a temperature rise was observed after a change from fertile to infertile mucus). This means that if a user stopped tracking before the Sympto algorithm closed the fertile window, their next period would not trigger the start of a new cycle, which would then lead to artificial ‘double cycles’.

Criteria summary for *standard cycle:*

- not the first cycle nor an on-going cycle

- observation gaps were no longer than 15 days

- at least one FAM body sign (BBT or cervical mucus or cervix position) was recorded

- no mid-cycle period-like bleeding detected when the cycle was longer than 40 days

- (Sympto only) defined as ovulatory cycles by the STM algorithm of Sympto, *i.e.* in which the fertile window could be closed.

- (Sympto only) no breastfeeding was reported or peri-menopause was declared

- (Kindara only) at least 8 FAM observations were reported

- (Kindara only) cycle length was at least 4 days longer than the total number of days in which bleeding was reported

***Cycles with reliable ovulation estimation*** (Sympto: 28,453 cycles; Kindara: 80,708)

Criteria summary for *cycles with reliable ovulation estimation*:

- *standard cycles*

- detected temperature shift was at least 0.15C or 0.27F. The temperature shift is estimated as the difference between the 25% percentile and the 80% percentile of temperature measurements of the current cycle. Temperature measurements are first given a weight (see below) and only measurements with a weight above 0.5 are considered for calculating this temperature shift.

- the uncertainty on the ovulation estimation was lower than ±1.5 day (see below for definition of uncertainty on estimation)

- the confidence score, which is related to acceptable amount of missing data in the ovulatory period, was equal or higher than 75% (see below for the definition of this confidence score)

**Selection criteria by figure**

Fig 2A, S1C: all users. However, axes were cut around “realistic values” such that typos or rare/extreme values like height of 50cm are excluded from these graphs.

Fig 2C, S1D, 3B-D, S2, S4A: all *standard cycles*.

Fig 4, S4B, S5, S6A-C: all *cycles with reliable ovulation estimation*.

Fig S6D: Sympto *cycles with reliable ovulation estimation* for which the uncertainty is lower than ½ day, the confidence is higher than 0.9 and the temperature shift higher than 0.35 C (429 cycles)

#### Observations decoding and ovulation timing estimation with HMM

The FAM body-signs are considered to reflect the hormonal changes orchestrating the menstrual cycles. Previous studies have shown that the combination of BBT and cervical mucus variations were reliable, although not perfect, proxies for the detection of ovulation (2, 3). Rise in BBT alone is not sufficient to detect ovulation in all women (4). Combining changes in mucus quality and quantity with BBT has been shown to improve the ovulation estimation (5), although this was not confirmed by other studies (6). Also, cervical mucus assessment is not an objective measure compared to BBT. We were interested in understanding which cycles were reported with observations that were consistent with previous literature description and for which it was thus possible to estimate timing of ovulation.

A diversity of methods can be considered for estimating the ovulation timing. We selected a Hidden Markov Model (HMM) as the most suitable due to its ability to uncover, from observations, latent phenomenon, which are here the cascade of hormonal events across the menstrual cycle. The choice of an HMM might, at first, seem counter-intuitive as transition probabilities could be expected to depend on the time already spent in a given state. However, we found that the model was robust to large variations in transition probabilities and that this framework relied on the user’s observations rather than on prior assumptions on cycle phase lengths to estimate ovulation time (Fig. S5, section “Robustness of ovulation estimation to model parameters”).

Briefly, we defined a 10-states HMM, in which each state is a particular phase of the menstrual cycle (Fig. 4A top, S3A, Methods), and used decoding algorithms to estimate the ovulation time, the uncertainty on this estimation, and a confidence score that accounts for missing observation and variation in temperature taking times.

We then established a set of stringent criteria on the uncertainty of the ovulation estimation (≤ ±1.5 days), the magnitude of the temperature shift (≥ 0.15 C) and the confidence score of the observations (≥ 0.75) to discriminate between cycles for which the estimations could be trusted (*cycles with reliable ovulation estimation*) and those where the observations did not allow for a reliable estimation of the ovulation day (Fig S4A, Methods). These strict criteria lead to the exclusion of ~40% (Sympto) and ~ 89% (Kindara) of the *standard cycles* that were initially selected. In total, 28,453 (Sympto) + 80,708 (Kindara) *cycles with reliable ovulation estimation* have been used for the subsequent analyses (Methods).

This mathematical framework for the estimation of the ovulation timing and of the different stages of the cycle holds some limitations. In particular, this estimation could benefit from the inclusion of non-ovulatory cycle detection and frequency assessment, thus generalizing the model to all cycles regardless of the occurrence of an ovulatory event. A framework providing a statistical probability of a cycle to be ovulatory or not could also be conceived, analogously to that proposed in (7) in the context of circadian rhythms.

**Model description**

The Hidden Markov Model (HMM) we defined describes a discretization in 10 states of the successive hormonal events throughout an ovulatory menstrual cycle. The HMM definition also includes the probabilities of observing the different FAM reported body signs in each state (emission probabilities) and the probabilities of switching from one state to another (transition probabilities). Emission probabilities were chosen to reflect observations previously made in studies that tested for ovulation with LH tests or ultrasounds (2, 6, 8), while transition probabilities were chosen in a quasi-uniform manner (see below).

Once the model was defined, we used the Viterbi and the Backward-Forward algorithms (9) to calculate the most probable state sequence for each cycle (Methods) and thus to have an estimation of ovulation timing, i.e. the most likely day of the cycle in which the HMM is in the state “ovulation”. We also computed the uncertainty of the estimation, *i.e.* the standard deviation of the distribution of probabilities for the state ‘ovulation’, which can be interpreted as the confidence interval in days for the time of ovulation estimation (see below). We also defined a confidence score that accounts for missing observations and variation in temperature taking time in a window of ~5 days around the estimated ovulation day (see below).

**HMM states**

We defined 10 states as a discretization of the hormonal evolution across the cycle.

*Menses*

HM: The first state (HM) is associated with the onset of the menses and the heavy/medium flow of fresh blood that characterize the first day(s) of menstruation.

LM: The second state (LM) is associated with the days of light bleeding or spotting that conclude menstruations.

*Follicular phase*

LE: When menses are over, the model switches to an early follicular state in which estrogen levels are still low (LE).

HE: As estrogen levels increase, and egg-white like or watery, stretchable cervical mucus might be produced and observed, the model evolves to a high estrogen (HE) state. Ovulation is expected to happen after a few days in that state (HE – late follicular phase) but, if ovulation fails, the model might stay in HE or switch back to LE, depending on the user’s observations.

*Ovulation*

Ovu: Ovulation being a brief event, the HMM ovulation state (Ovu) is the only state in which the model can only stay for one day before transitioning to one of the two possible early luteal phase states.

*Luteal phase*

Luteal phase is marked by the progressive production of progesterone by the corpus luteum.

Rise/HP: Progesterone rise might be fast or slow and thus, after ovulation, the model can switch and stay for some days in a ‘Rise’ state or switch straight to a high progesterone (HP – early luteal phase) stage.

EP: After a drop following ovulation, estrogens usually increase again due to production by the corpus luteum, and this estrogen peak is sometimes observed by users via a small cervical mucus production, which led us to define a “EP” state for the possible observation of higher estrogen levels.

LP: The decrease in hormones in the late luteal phase announces the end of the cycle and the expected next menses. In that state (LP for low progesterone), no mucus is usually observed, temperature might drop slightly, and light bleeding can be observed.

*End*

We added an artificial observation at the end of each cycle, which can only be observed with probability 1 in an extra “end” state. This state ensures that the model properly evolves from menses through ovulation until the last observations for each *standard cycle*.

**Transition probabilities**

As we wanted the HMM decoding to be mostly influenced by the observations and not so much by prior hypotheses on the transitions from one state to another to reflect the diversity of menstrual patterns between individuals and cycles, we chose quasi-uniform probabilities, e.g., if one can switch to n states from an initial one, then the transition probabilities were set to 1/n. Exceptions to that rule were made for transitions from “Rise”, “HP” and “EP”.

Transition probabilities are given in Table S1. Initial probabilities were set to 1 on the “HM” state.

State definition and transition probabilities were set to the same values for Kindara and Sympto users.

**Emission probabilities**

Emission probabilities reflect the probability of observing a set of given body signs in each state of the cycle. Given that the datasets do not have positive controls (e.g. clinically confirmed ovulation or hormonal levels) to train these emission probabilities, we defined them based on existing literature.

For each of the FAM observations, we also included a probability for that observation to not be reported which changes over the cycle. These emission probabilities were set differently for Sympto and Kindara datasets to reflect the slight variations in their tracking options.

*Bleeding*

The emission probabilities for the bleeding observation directly follows the definition of the menses states and observations in Fig 2. Heavy bleeding is most commonly found at the beginning of the menses, in the state HM while lighter bleeding follows (in the state LM). As we noticed that light bleeding was also reported by the end of the cycle, especially when temperature was dropping (Fig 3), we increased the probability to observe light bleeding in the ‘LP’ state. Possibility of ovulation bleeding is also included in the bleeding emission matrices. See Tables S2-3.

*Temperature emission probabilities*

Based on previous results (2, 5, 6, 10), and on the observations obtained in Fig 2, we defined the temperature emission probabilities as follow :

For each state, we draw a normal distribution around the expected temperature in that menstrual state: typically, low for follicular states and high for luteal states. The low temperature (T.low) is calculated as the 25 percentile, with the high temperature (T.high) is calculated at the 80 percentile. The temperature shift is calculated as the difference between these two temperatures. The normal distributions (denoted as N(mean, sd) in Table S4) are normalized so that their sum over the temperature range is equal to 1. We then define the probabilities of not observing any temperature, which is higher at the beginning and the end of the cycle, or to observe extreme temperature that fall outside the defined temperature range. The whole matrix as presented in Table S4 is then normalized so that the sum of probabilities by state is equal to 1.

*Cervical mucus*

Cervical mucus production has been shown to vary in quantity and quality over the menstrual cycle. Its biophysical properties, such as its viscosity, composition or “Spinbarkeit” have been extensively described (6, 8, 11–14). When app users observe their cervical mucus, they usually do this by visual inspection and by stretching the mucus between their fingers. By looking at the color (is the mucus transparent or translucent?), the stretchiness (can the mucus be stretched over 2, 3 cm?), its consistency (is the mucus thick or thin?), they classify their mucus into different categories. Kindara offers a larger diversity of options for the mucus classification (Table 2) and the emission probabilities reflect that larger diversity. The probabilities have been establish in order to reflect the previous findings that emerged from cohorts in which the hormonal profiles or events had been clinically assessed (2, 6).

*Cervix position and vaginal sensation*

Similarly, the cervix has been shown to exhibit physical changes across the menstrual cycles (Fig S2).

We used these observations to establish the emission probabilities found in Table S9 and S10. Here again, Kindara offers broader options to track the cervical characteristics by letting users track independently its height, openness and firmness. In order to simplify the model, we combined these 3 characteristics into one, by taking the observation that showed the highest probability of fertility. For example, if cervical height is 1, cervical openness is 2 and cervical firmness is 1, we will attribute a score of 2 (max of the 3 observations). Sympto presents users with only 3 options (Table 2).

Vaginal sensation has also been reported to vary during the menstrual cycle, with the vagina feeling the most wet or lubricated in the days leading up to ovulation. Due to the extremely small fraction of Kindara users reporting their vaginal sensation, we did not use that variable. Sympto offer their users to either track cervical position or their vaginal sensation according to what they think is easier to track. We thus have an emission matrix that combines cervical position and vaginal feel for Sympto (Table S5-8)

**Estimation of ovulation day**

To estimate the most likely day of ovulation, we used two algorithms available in the R package hmm (9). First, we used the Viterbi algorithm. This algorithm provides the most likely sequence of hidden state given the observations.

We then used the Backward-Forward algorithm (posterior function from the same R package). This algorithm provides the probabilities at each day of the cycle to be in a given state. If we thus look at the probabilities of the state ‘Ovu’ (ovulation day) for each day of the cycle, we can assess on which day that probability was the highest. We can also look at the distribution of these probabilities and estimate ovulation as the weighted average of these probabilities. For example, if the probability to be in state “Ovu” is 1 on day 14, we can consider that the most likely day of ovulation is day 14. However, if the probability of state “Ovu” is 0.5 on both days 14 and 15, we could use the weighted sum and provide an estimate of 14.5 for the timing of ovulation.

In Fig 3,4 and S3 the line labeled “hmm” shows the most likely sequence of hidden states as computed with the Viterbi algorithm, while the line plot below that line shows the probability of each hidden states for each cycle day.

The most likely day of ovulation (*O*) is calculated as $O=d . p_{d}^{O}$ where $d$ is the cycleday and $p_{d}^{O}$ is the probability to be in state “Ovu” on cycle day $d$.

**Uncertainty on the ovulation estimation**

The posterior probabilities provided by the Backward-Forward algorithm were also used to define a measure of uncertainty on the estimation of the ovulation day. Indeed, if probabilities for the state « Ovu » peak at 1 on a given day, we have a high confidence on the estimation of ovulation timing. On the contrary, if the probabilities are distributed over many days, we can infer that the uncertainty on the ovulation estimate is high.

We computed this uncertainty (*u*) as $u= \sqrt{p_{d}^{O}.{(O-d)}^{2}}$ and can be interpreted as the standard deviation of the « Ovu » state probabilities distribution.

**Confidence score**

We further defined a confidence score that reflects the amount and trust in observed data that was available around the estimated timing of ovulation. For example, we have a higher confidence in the ovulation estimation if temperature has been recorded every day at the same time and if cervical mucus has been reported every day. On the other side, if few observations have been reported, we have a lower confidence in the ovulation estimation.

The confidence score $C$ is defined as follow: $C= \left( C^{T}+ C^{M}+C^{w_{T}} \right)/3$

Where $C^{T}$ and $C^{M}$reflect the number of temperature ($T$) and mucus ($M$) observations reported.

They are calculated as follow:

$$C^{T/M}=\sum_{d= ovu-3}^{ovu+3} N(\mu= 0, \sigma= 2) *B_{d}(measurement)$$

Where $B_{d}$ is a Boolean that is equal to 1 if a measurement is reported and to 0 if the measurement is missing. *N* denotes a normal distribution.

$C^{w_{T}}$ reflects the reliability of the temperature measurement. It is the average of the temperature weights (below).

**Weights on BTT measurements**

Weights on each temperature data point were defined to take into account the regularity of temperature measurement times, to give less weight to measurements at the very beginning of the cycle (temperature has been observed to change during the menses) and to attenuate outlier contributions.

Weights were computed as follow:

$$w_{d}=\left( 1- {pT}_{d} \right)\left( {1- pB}_{d} \right) \left( 1-{pO}_{d} \right)$$

where $w_{d}$ is the weight applied to temperature measurement at day $d$, ${pT}_{d}$ is the penalty associated to deviation of the time at which basal body temperature was taken, ${pB}_{d}$ is the penalty for measurements at the beginning of the cycle and ${pO}_{d}$ is the penalty for outliers. Their mathematical expression is the following:

$${pT}_{d}=\frac{0.5}{1+e^{-\frac{2C}{1.8}\left( \left| T_{d}-T_{M} \right|-1 \right)}}$$

where $C=log\left( {0.95}/{0.05} \right)$, $T_{d}$ is temperature measured at day *d* and $T_{M}$ is the median temperature over the cycle.

$${pB}_{d}=0.5 if d<6, 0.25 if d=6, 0 if d>6$$

${pO}_{d}=1- \frac{{diff}_{d}}{2 dt}$

where ${diff}_{d}$ is the maximum of the absolute difference between a measurement and its neighboring measurement and $dt$ is the maximum between 1F or 0.5 C and the max of ${diff}_{d}$ for all temperature measurements in that cycle.

**Robustness of ovulation estimation to model parameters and missing data**

*Transition probabilities*

To test whether we could confidently use quasi-uniform transition probabilities, we ran the HMM decoding on a subset of the data and increasingly added random noise to the transition probabilities such as

$$q_{x,i}= \frac{p_{x,i} {10}^{\varepsilon_{i}}}{\sum_{j=1}^{N} p_{x,j}{10}^{\varepsilon_{j}}}$$

where $q_{x,i}$ is the noisy transition probability from a given state $x$ to a state $i$ , $p_{x,i}$ is the original (quasi-uniform) transition probability and $\varepsilon_{i}$ is a random variable drawn from a normal distribution $\varepsilon\sim N(0,\sigma)$.

We observed that, even for quasi random probabilities (when $\sigma$ is large), the difference between the estimated ovulations with uniform or random probabilities stays small and lower or in the range of the estimation uncertainties (Fig S6A).

Similarly, instead of random noise, we increasingly diagonalized the transition probability matrix by increasing the probability of staying in a given state:

$$q_{x,i}= \frac{r_{x,i}}{\sum_{j=1}^{N} r_{x,j}} ;r_{x,i}=p_{x,i}+ p_{x,i}*C \text{if }x= i; r_{x,i}=p_{x,i}\text{ otherwise}$$

Shifting away from quasi-uniform transition probabilities (larger values of $C$) only slightly impacted the ovulation estimation, with a small trend towards ovulation estimated later in the cycle (shorter luteal phases) (Fig S6B).

*Emission probabilities*

In comparison, random noise on the emission probabilities have a much higher impact on the ovulation estimation as seen on Fig S6C. In this case, emission probabilities have been modified as follow:

$$q_{x,i}= \frac{p_{x,i} {10}^{\varepsilon_{i}}}{\sum_{j=1}^{N} p_{x,j}{10}^{\varepsilon_{j}}}$$

where $q_{x,i}$ is the noisy emission probability of emitting observation set $i$ from a given state $x$, $p_{x,i}$ is the original emission probability (based on existing literature as described above) and $\varepsilon_{i}$ is a random variable drawn from a normal distribution $\varepsilon\sim N(0,\sigma)$.

*Missing data*

To investigate the impact of missing temperature or cervical mucus reports on ovulation estimation, profiles, who had low uncertainty and high estimation confidence were selected and records were progressively removed from these profiles. Ovulation time was then estimated along with the estimation uncertainty and confidence using the same method as previously described. Results show that missing temperature records tends to slightly advance ovulation time estimation (by ~6h on average if all temperature records are ignored for the estimation). Missing temperature records leads to larger estimation uncertainties than missing cervical mucus reports. These results suggest that temperature is slightly more useful than cervical mucus to estimate the ovulation time (Fig S6D).

### Supplementary Figures

**Fig. S1**

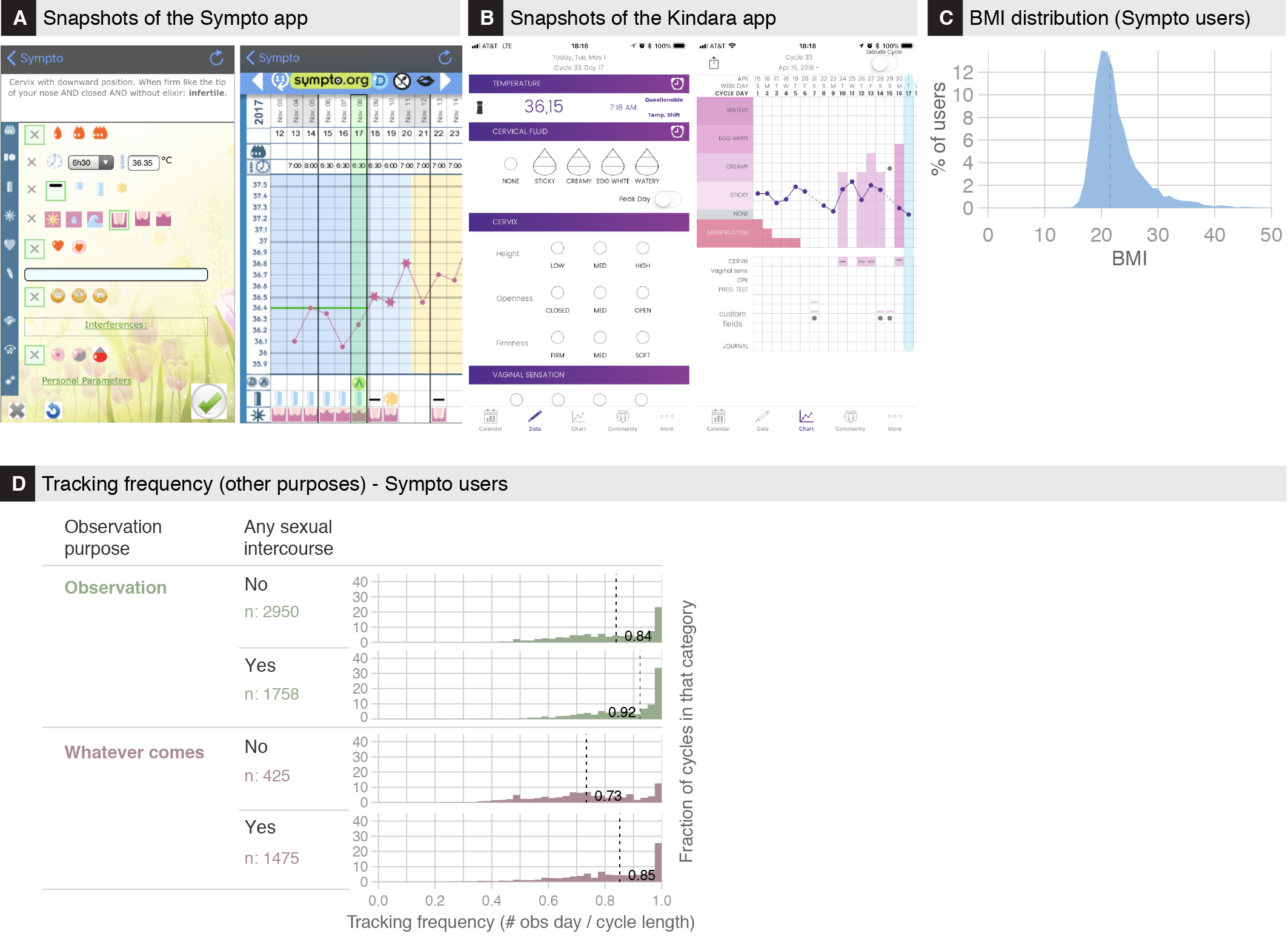

**Figure S1**
**(A)** Screenshots from the Sympto app as displayed to iPhone users. (Left) Observation tracking dashboard. (Right) “Cyclogramme”: visual display of reported observations.

**(B)** Screenshots from the Kindara app as displayed to iPhone users. (Left) Observation tracking dashboard. (Right) Visual display of reported observations.

**(C)** BMI (kg/m^2^) distribution of Sympto users.

**(D)** Tracking frequency (number of days with at least one observation / number of days in this cycle) of cycles for which the tracking purpose has been declared as “Observation” or “Accept what comes” by Sympto users at the beginning of the cycle.

**Fig. S2**

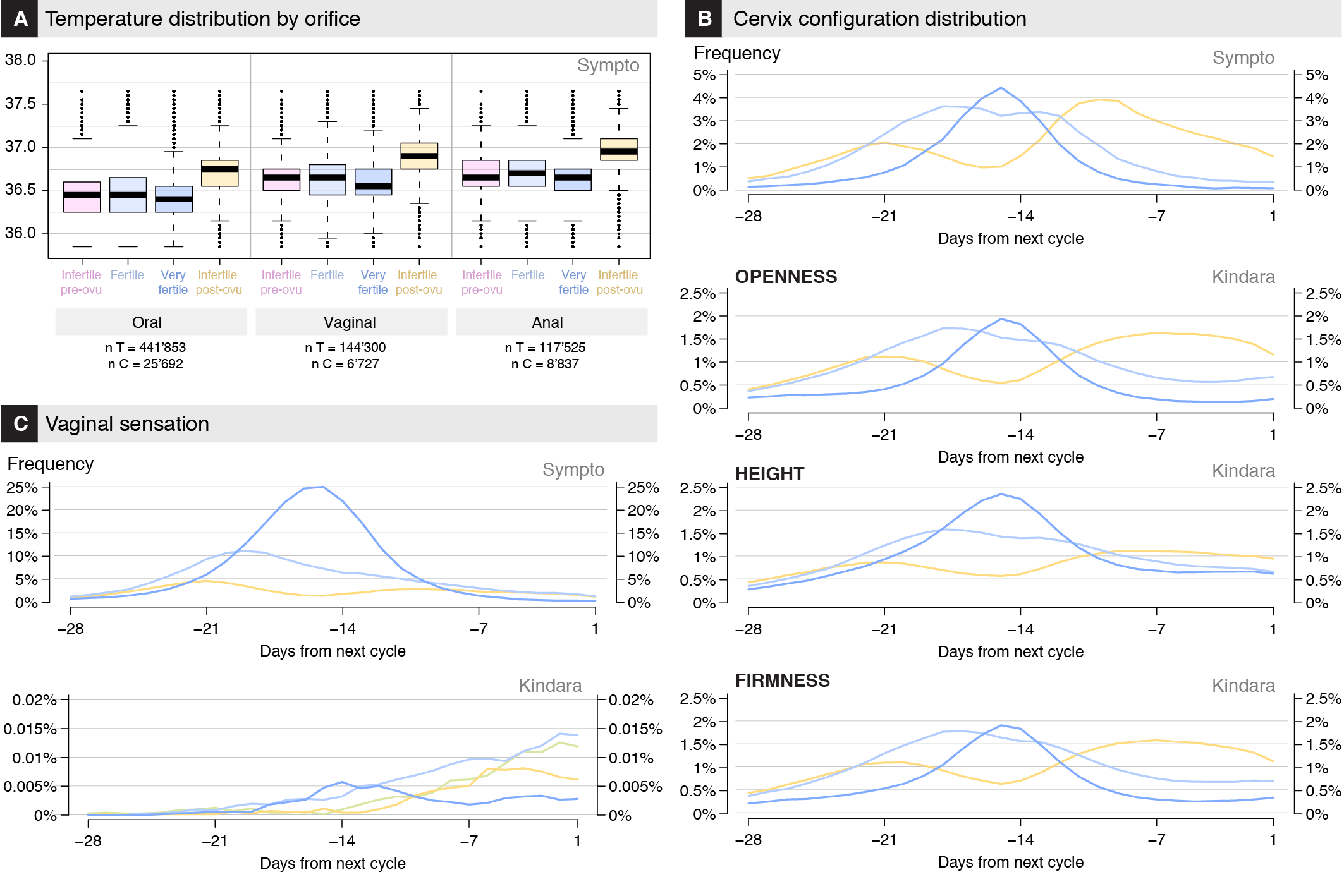

**Figure S2**
**(A)** Temperature distribution by orifice for Sympto users that are prompted to choose at the beginning of the cycle and to keep that option for the full duration of the cycle. The different phases (infertile pre-ovu, fertile, very fertile or infertile post-ovu) are the phases as calculated by the Sympto algorithm.

**(B)** Frequency of cervix observations. For Sympto, yellow indicates “low/closed/firm”, dark blue indicated “high/open/soft” while light blue reflects an intermediary configuration. For Kindara, yellow indicates low (HEIGHT) or closed (OPENNESS) or firm (FIRMNESS), dark blue indicates high (HEIGHT) or open (OPENNESS) or soft (FIRMNESS) while light blue reflects intermediary levels.

**(C)** Frequency of vaginal sensation. (Sympto) yellow: dry, light blue: little moist, dark blue: very wet, moist, lubricated. (Kindara) yellow: dry-sticky, beige-green: dry, light blue: wet moist, dark blue: wet lubricative.

**Fig. S3**

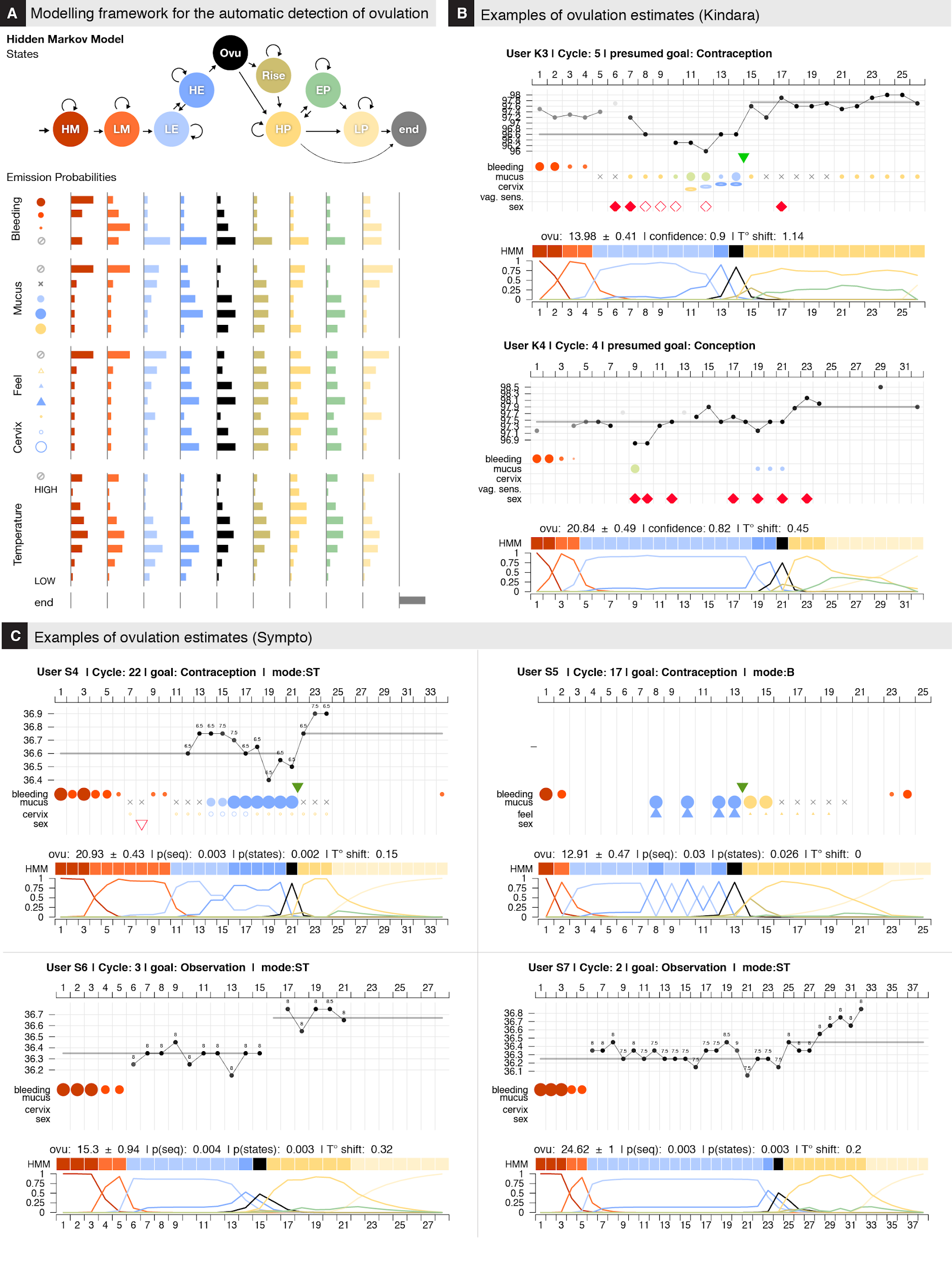

**Figure S3**
**(A)** Illustration of the modeling framework used for the estimation of ovulation timing. (Top) Schematic representation of the HMM with the 10 states that have been defined to discretize the menstrual hormonal events. Full description of the HMM setup is provided in the Methods. In short, ‘HM’ stands for ‘heavy Menses’, ‘LM’ for ‘light Menses’, ‘LE’ for ‘low Estrogen’, ‘HE’ for ‘high Estrogen’, ‘Ovu’ for ‘Ovulation’, ‘Rise’ for ‘progesterone/BTT rise’, ‘HP’ for ‘high Progesterone’,’EP’ for ‘Estrogen peak in luteal phase’, ‘LP’ for ‘low Progesterone’. The arrows between the states indicate possible transition; arrow thickness is not representative of transition probabilities. Values of the transition probability matrix can be found in the Methods. (Bottom) Emission probabilities for the FAM body signs other than BBT. Exact values and supporting references can be found in the Methods.

**(B)** Example of results from the menstrual state estimation framework for the 2rd and 3rd cycle of 2 different Kindara users. (Top of each chart) Original user observations as in Fig. 2A. (Middle of each chart) The square colors in the HMM-labeled line represent the most likely sequence of HMM states given the observations in the cycle as calculated with the Viterbi algorithm (‘viterbi’ function in R, ‘HMM’ package)(9). (Bottom of each chart) Normalized probabilities of each state on each day of the cycle, as calculated with the posterior algorithm (‘posterior’ function in R, ‘HMM’ package).

**(C)** Idem as (B) but for cycles of 4 different Sympto users.

**Fig. S4**

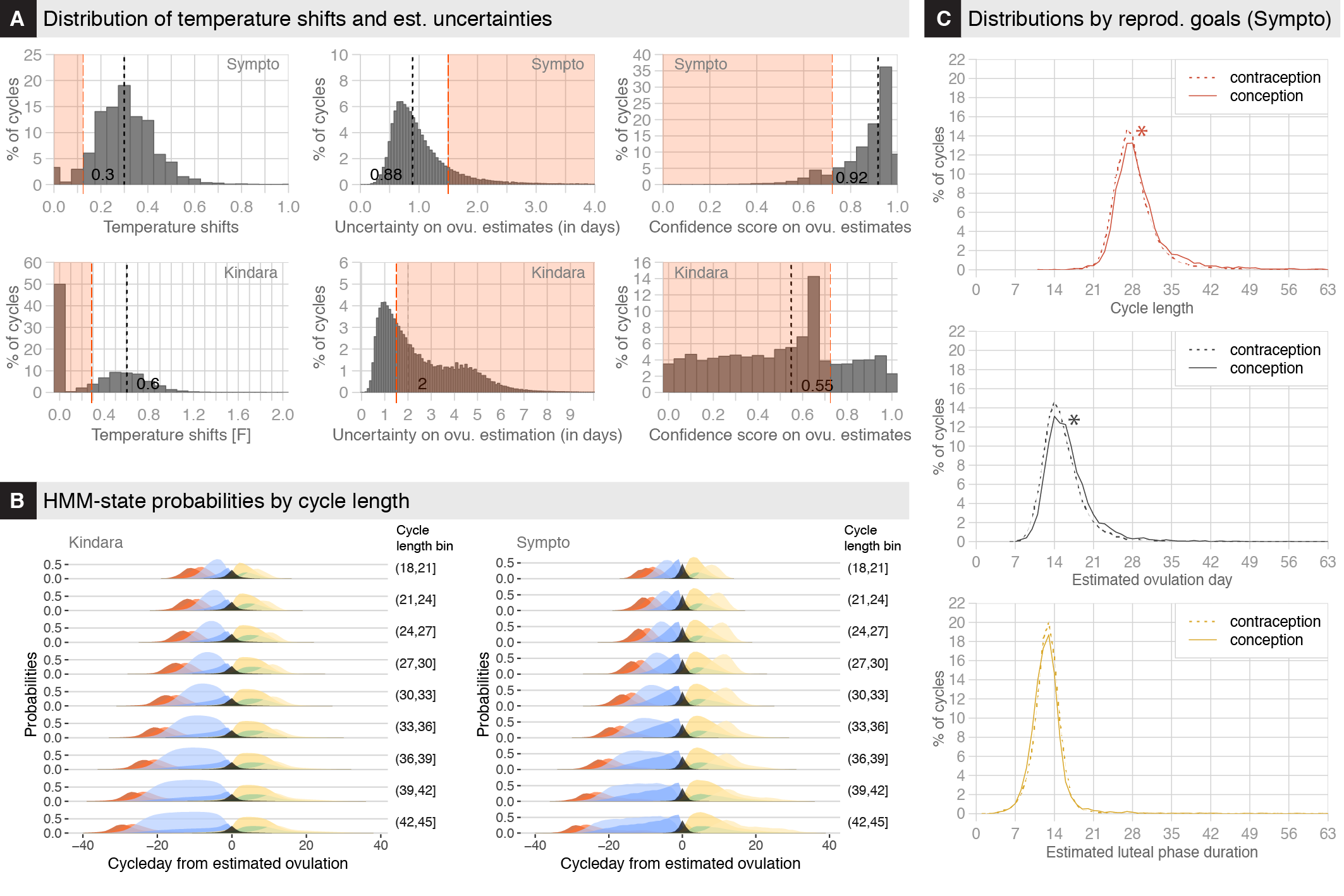

**Figure S4**
**(A)** Distribution of detected temperature shifts (Left), ovulation estimation uncertainties (Middle) and confidence score (Right). We observe similar temperature shift distribution for cycles logged in Kindara or Sympto, but we find a lot more cycles in which temperature has not been logged at all or less than 4 time in cycles reported via the Kindara app. Temperature shift are calculated as the difference between the 25 and 80 percentiles of the distribution of acceptable temperature for each cycle (Methods). Because of these missing temperature recordings, the uncertainty distribution on Kindara cycles also show a much heavier tail on higher values. This heavier tail might also reflect observations that are not coherent with the current knowledge of body sign variation in respect to the menstrual cycle, which are less likely to be observed in Sympto cycles as users are more likely to receive human counselling on the interpretation of their observations. Similarly, the confidence score, which reflects missing observations around the estimated ovulation time, show a higher distribution in Kindara cycles.

**(B)** Average estimated state probabilities by cycle-day counting from estimated ovulation aggregated by total cycle length (in bins of 3 units) for all *cycles with reliable ovulation estimation* separately for Kindata (left) and Sympto (right)

**(C)** Distribution of cycle length, estimated ovulation day and estimated luteal phase duration for Sympto cycles according to the declared goals (avoiding or seeking pregnancy) for that specific cycle. We observe significantly different distributions (p<2.2-16) for the cycle length and ovulation time (p = 0.08 for luteal phase). These differences might reflect the fact that users seeking pregnancy might be at higher risks of subfertility if we assume that they could not conceive for some month before deciding using the app. We controlled for the age of users by running a logistic regression and found that the goals explained more of the variance than the age (cycle_length ~ 34.3 - 0.15 x age_at_obs + 0.57 x goal_LR -0.03 x cycle_nb; ovu ~ 21.22 -0.14 x age_at_obs + 0.44 x goal_LR + -0.04 cycle_nb)

**Fig. S5**

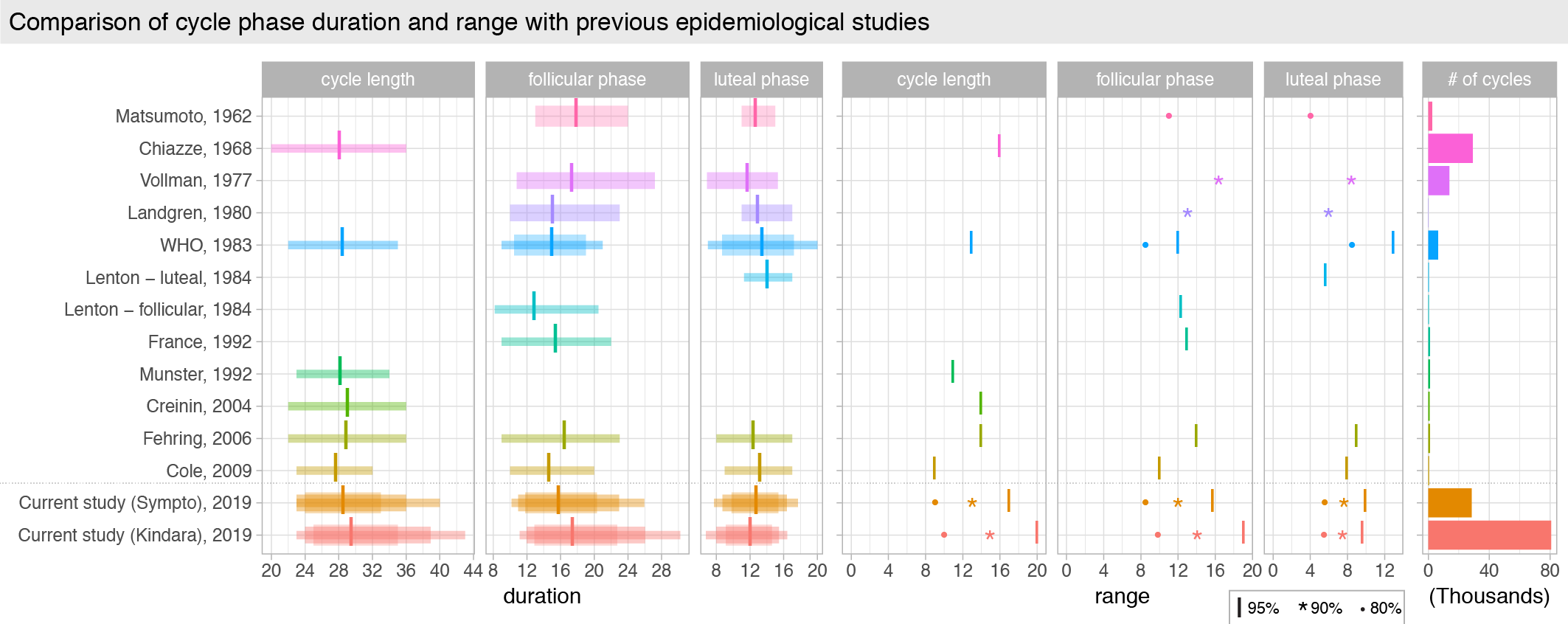

**Figure S5**

Comparison with previous epidemiological studies of the cycle length, follicular phase and luteal phase duration and ranges, in days, as well as the number of cycles included for each of these studies.

**Fig. S6**

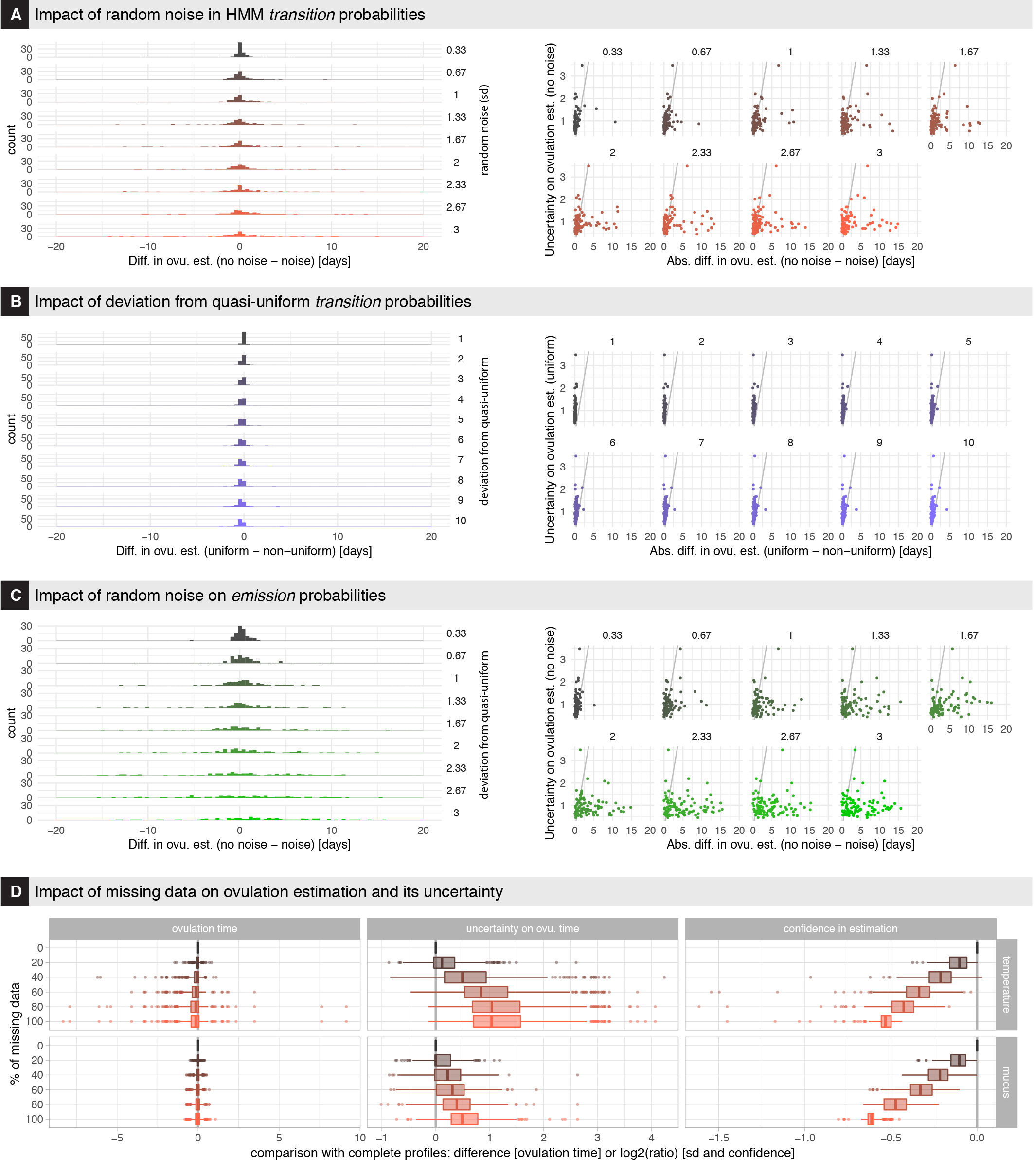

**Figure S6**
**(A-C)** Distribution of differences in ovulation estimation (Left). Ovulation uncertainties vs absolute difference in ovulation estimation (Right).

**(A)** Random noise was added to the transition probabilities.

**(B)** Self-transition probabilities were increased.

**(C)** Random noise was added to the emission probabilities.

**(D)** Impact of missing temperature or cervical mucus records on ovulation estimation and on the estimation uncertainty and confidence.

### Supplementary tables

#### Table S1: transition probabilities

| from\to | hM | lM | lE | hE | O | Rise | hP | Ep | lP | end |
| --- | --- | --- | --- | --- | --- | --- | --- | --- | --- | --- |
| hM | 0.5 | 0.5 | 0 | 0 | 0 | 0 | 0 | 0 | 0 | 0 |
| lM | 0 | 0.5 | 0.5 | 0 | 0 | 0 | 0 | 0 | 0 | 0 |
| lE | 0 | 0 | 0.5 | 0.5 | 0 | 0 | 0 | 0 | 0 | 0 |
| hE | 0 | 0 | 1/3 | 1/3 | 1/3 | 0 | 0 | 0 | 0 | 0 |
| O | 0 | 0 | 0 | 0 | 0 | 0.5 | 0.5 | 0 | 0 | 0 |
| Rise | 0 | 0 | 0 | 0 | 0 | 0.4 | 0.6 | 0 | 0 | 0 |
| hP | 0 | 0 | 0 | 0 | 0 | 0 | 0.55 | 0.2 | 0.2 | 0.05 |
| Ep | 0 | 0 | 0 | 0 | 0 | 0 | 0.35 | 0.3 | 0.35 | 0 |
| lP | 0 | 0 | 0 | 0 | 0 | 0 | 0 | 0 | 0.5 | 0.5 |
| end | 0 | 0 | 0 | 0 | 0 | 0 | 0 | 0 | 0 | 1 |

#### Table S2: Emission probabilities bleeding – Kindara

| state\bleeding | NA | 3 | 2 | 1 | s | end |
| --- | --- | --- | --- | --- | --- | --- |
| hM | 0.141 | 0.423 | 0.423 | 0.007 | 0.007 | 0 |
| lM | 0.146 | 0.008 | 0.308 | 0.308 | 0.231 | 0 |
| lE | 0.773 | 0.045 | 0.045 | 0.045 | 0.091 | 0 |
| hE | 0.773 | 0.045 | 0.045 | 0.045 | 0.091 | 0 |
| O | 0.667 | 0.048 | 0.048 | 0.095 | 0.143 | 0 |
| Rise | 0.701 | 0.047 | 0.047 | 0.065 | 0.14 | 0 |
| hP | 0.81 | 0.048 | 0.048 | 0.048 | 0.048 | 0 |
| Ep | 0.81 | 0.048 | 0.048 | 0.048 | 0.048 | 0 |
| lP | 0.5 | 0.042 | 0.042 | 0.167 | 0.25 | 0 |
| end | 0 | 0 | 0 | 0 | 0 | 1 |

#### Table S3: Emission probabilities bleeding – Sympto

| state\bleeding | NA | 3 | 2 | 1 | end |
| --- | --- | --- | --- | --- | --- |
| hM | 0.2 | 0.6 | 0.19 | 0.01 | 0 |
| lM | 0.19 | 0.01 | 0.4 | 0.4 | 0 |
| lE | 0.85 | 0.05 | 0.05 | 0.05 | 0 |
| hE | 0.85 | 0.05 | 0.05 | 0.05 | 0 |
| O | 0.7 | 0.05 | 0.05 | 0.2 | 0 |
| Rise | 0.75 | 0.05 | 0.05 | 0.15 | 0 |
| hP | 0.85 | 0.05 | 0.05 | 0.05 | 0 |
| Ep | 0.85 | 0.05 | 0.05 | 0.05 | 0 |
| lP | 0.6 | 0.05 | 0.05 | 0.3 | 0 |
| end | 0 | 0 | 0 | 0 | 1 |

#### Table S4: Emission probabilities for temperature - Sympto & Kindara

| state\BBT | NA | LOW | T.low-DT ….. T.high+DT | HIGH | end |
| --- | --- | --- | --- | --- | --- |
| hM | 0.5 | 0.17 | N(mean = T.low + DT/3 , sd = DT/2 ) | 0.13 | 0 |
| lM | 0.4 | 0.13 | N(mean = T.low + DT/4 , sd = DT/2 ) | 0.1 | 0 |
| lE | 0.2 | 0.1 | N(mean = T.low , sd = DT/2 ) | 0.07 | 0 |
| hE | 0.1 | 0.07 | N(mean = T.low , sd = DT/2 ) | 0.03 | 0 |
| O | 0.1 | 0.03 | N(mean = T.low + DT/4, sd = DT/2 ) | 0.07 | 0 |
| Rise | 0.1 | 0.03 | N(mean = T.high- DT/3 , sd = DT/2 ) | 0.07 | 0 |
| hP | 0.2 | 0.03 | N(mean = T.high , sd = DT/2 ) | 0.1 | 0 |
| Ep | 0.3 | 0.03 | N(mean = T.high- DT/3 , sd = DT/2 ) | 0.1 | 0 |
| lP | 0.4 | 0.1 | N(mean = T.low + DT/3, sd = DT/2 ) | 0.07 | 0 |
| end | 0 | 0 | 0 | 0 | 1 |

#### Table S5: Emission probabilities cervical mucus – Kindara

| state\mucus | NA | No | s1 | s2 | s3 | c1 | c2 | c3 | e1 | e2 | e3 | w1 | w2 | w3 | end |
| --- | --- | --- | --- | --- | --- | --- | --- | --- | --- | --- | --- | --- | --- | --- | --- |
| hM | 0.333 | 0.267 | 0.033 | 0.033 | 0.033 | 0.033 | 0.033 | 0.033 | 0.033 | 0.033 | 0.033 | 0.033 | 0.033 | 0.033 | 0 |
| lM | 0.333 | 0.267 | 0.033 | 0.033 | 0.033 | 0.033 | 0.033 | 0.033 | 0.033 | 0.033 | 0.033 | 0.033 | 0.033 | 0.033 | 0 |
| lE | 0.182 | 0.182 | 0.073 | 0.036 | 0.018 | 0.145 | 0.127 | 0.091 | 0.036 | 0.018 | 0.018 | 0.036 | 0.018 | 0.018 | 0 |
| hE | 0.043 | 0.043 | 0.014 | 0.014 | 0.014 | 0.029 | 0.043 | 0.058 | 0.116 | 0.13 | 0.145 | 0.116 | 0.116 | 0.116 | 0 |
| O | 0.034 | 0.034 | 0.056 | 0.067 | 0.079 | 0.011 | 0.022 | 0.022 | 0.112 | 0.112 | 0.112 | 0.112 | 0.112 | 0.112 | 0 |
| Rise | 0.043 | 0.071 | 0.1 | 0.1 | 0.1 | 0.029 | 0.029 | 0.043 | 0.1 | 0.1 | 0.086 | 0.086 | 0.071 | 0.043 | 0 |
| hP | 0.14 | 0.14 | 0.14 | 0.105 | 0.053 | 0.105 | 0.123 | 0.088 | 0.018 | 0.018 | 0.018 | 0.018 | 0.018 | 0.018 | 0 |
| Ep | 0.125 | 0.125 | 0.167 | 0.062 | 0.021 | 0.104 | 0.104 | 0.104 | 0.042 | 0.042 | 0.021 | 0.042 | 0.021 | 0.021 | 0 |
| lP | 0.244 | 0.244 | 0.073 | 0.049 | 0.024 | 0.098 | 0.073 | 0.049 | 0.024 | 0.024 | 0.024 | 0.024 | 0.024 | 0.024 | 0 |
| end | 0 | 0 | 0 | 0 | 0 | 0 | 0 | 0 | 0 | 0 | 0 | 0 | 0 | 0 | 1 |

#### Table S6: Emission probabilities cervical mucus – Sympto

| state\mucus | NA | No | f | F | s | end |
| --- | --- | --- | --- | --- | --- | --- |
| hM | 0.45 | 0.4 | 0.05 | 0.05 | 0.05 | 0 |
| lM | 0.45 | 0.4 | 0.05 | 0.05 | 0.05 | 0 |
| lE | 0.35 | 0.35 | 0.25 | 0.05 | 0.05 | 0 |
| hE | 0.1 | 0.1 | 0.35 | 0.4 | 0.05 | 0 |
| O | 0.1 | 0.1 | 0.15 | 0.5 | 0.15 | 0 |
| Rise | 0.1 | 0.1 | 0.3 | 0.2 | 0.3 | 0 |
| hP | 0.3 | 0.3 | 0.1 | 0.1 | 0.2 | 0 |
| Ep | 0.1 | 0.1 | 0.3 | 0.2 | 0.3 | 0 |
| lP | 0.4 | 0.4 | 0.05 | 0.05 | 0.1 | 0 |
| end | 0 | 0 | 0 | 0 | 0 | 1 |

#### Table S7: Emission probabilities cervix position – Kindara

| state\cervix | NA | 1 | 2 | 3 | end |
| --- | --- | --- | --- | --- | --- |
| hM | 0.714 | 0.071 | 0.143 | 0.071 | 0 |
| lM | 0.588 | 0.118 | 0.176 | 0.118 | 0 |
| lE | 0.588 | 0.118 | 0.176 | 0.118 | 0 |
| hE | 0.333 | 0.1 | 0.267 | 0.3 | 0 |
| O | 0.345 | 0.069 | 0.241 | 0.345 | 0 |
| Rise | 0.357 | 0.143 | 0.286 | 0.214 | 0 |
| hP | 0.476 | 0.286 | 0.143 | 0.095 | 0 |
| Ep | 0.588 | 0.118 | 0.176 | 0.118 | 0 |
| lP | 0.455 | 0.136 | 0.182 | 0.227 | 0 |
| end | 0 | 0 | 0 | 0 | 1 |

#### Table S8: Emission probabilities cervix position and vaginal sensation – Sympto

| state\cervix | NA | FD | FW | FVW | CC | CM | CH | end |
| --- | --- | --- | --- | --- | --- | --- | --- | --- |
| hM | 0.143 | 0.143 | 0.143 | 0.143 | 0.143 | 0.143 | 0.143 | 0 |
| lM | 0.143 | 0.143 | 0.143 | 0.143 | 0.143 | 0.143 | 0.143 | 0 |
| lE | 0.192 | 0.192 | 0.192 | 0.019 | 0.192 | 0.192 | 0.019 | 0 |
| hE | 0.143 | 0.071 | 0.143 | 0.214 | 0.071 | 0.143 | 0.214 | 0 |
| O | 0.217 | 0.022 | 0.109 | 0.217 | 0.043 | 0.174 | 0.217 | 0 |
| Rise | 0.143 | 0.143 | 0.143 | 0.143 | 0.143 | 0.143 | 0.143 | 0 |
| hP | 0.278 | 0.278 | 0.056 | 0.028 | 0.278 | 0.056 | 0.028 | 0 |
| Ep | 0.417 | 0.083 | 0.167 | 0.042 | 0.083 | 0.167 | 0.042 | 0 |
| lP | 0.312 | 0.25 | 0.062 | 0.031 | 0.25 | 0.062 | 0.031 | 0 |
| end | 0 | 0 | 0 | 0 | 0 | 0 | 0 | 1 |

#### Table S9: Cycle phases duration and ranges of previous studies

| year | study | method | exclusion | n individuals | n cycles | phase | mean | SD | perc 2.5 | perc 5 | perc 10 | perc 90 | perc 95 | perc 97.5 |
| --- | --- | --- | --- | --- | --- | --- | --- | --- | --- | --- | --- | --- | --- | --- |
| 2009 | Cole | daily urine LH HCG | anovulatory or long cycles | 167 | 458 | cycle length | 27.7 | 2.4 | 23 |  |  |  |  | 32 |
| 2009 | Cole | daily urine LH HCG | anovulatory or long cycles | 167 | 458 | ovulation | 14.7 | 2.4 | 10 |  |  |  |  | 20 |
| 2009 | Cole | daily urine LH HCG | anovulatory or long cycles | 167 | 458 | luteal | 13.2 | 2 | 9 |  |  |  |  | 17 |
| 1984 | Lenton - follicular | blood LH | anovulatory or long cycles | 293 | 327 | ovulation | 12.94 |  | 8.2 |  |  |  |  | 20.5 |
| 1984 | Lenton - luteal | blood LH | anovulatory or long cycles | 293 | 327 | luteal | 14.13 | 1.41 | 11.3 |  |  |  |  | 17 |
| 1983 | WHO | mucus | long cycles | 687 | 6427 | cycle length | 28.5 | 3.2 | 22 |  |  |  |  | 35 |
| 1983 | WHO | mucus | long cycles | 687 | 6427 | ovulation | 15 | 2.6 | 9 |  | 10.5 | 19 |  | 21 |
| 1983 | WHO | mucus | long cycles | 687 | 6427 | luteal | 13.5 | 2.8 | 7 |  | 8.7 | 17.2 |  | 20 |
| 1980 | Landgren | blood LH | long cycles | 68 | 68 | ovulation | 15.1 |  |  | 10 |  |  | 23 |  |
| 1980 | Landgren | blood LH | long cycles | 68 | 68 | luteal | 13 |  |  | 11 |  |  | 17 |  |
| 1977 | Vollman | BBT | non-biphasic | 524 | 13785 | ovulation | 17.4 |  |  | 10.8 |  |  | 27.2 |  |
| 1977 | Vollman | BBT | non-biphasic | 524 | 13785 | luteal | 11.7 |  |  | 6.9 |  |  | 15.3 |  |
| 1962 | Matsumoto | BBT | non-biphasic |  | 2500 | ovulation | 17.9 |  |  |  | 13 | 24 |  |  |
| 1962 | Matsumoto | BBT | non-biphasic |  | 2500 | luteal | 12.7 |  |  |  | 11 | 15 |  |  |
| 2006 | Fehring | fertility monitor | | 141 | 1060 | cycle length | 28.9 | 3.4 | 22 |  |  |  |  | 36 |
| 2006 | Fehring | fertility monitor | | 141 | 1060 | ovulation | 16.5 | 3.4 | 9 |  |  |  |  | 23 |
| 2006 | Fehring | fertility monitor | | 141 | 1060 | luteal | 12.4 | 2 | 8 |  |  |  |  | 17 |
| 1968 | Chiazze |  |  | 2316 | 29238 | cycle length | 28.1 | 4 | 20 |  |  |  |  | 36 |
| 2004 | Creinin |  |  | 130 | 786 | cycle length | 29.1 | 3.5 | 22 |  |  |  |  | 36 |
| 1992 | Munster |  |  | 1128 | 1128 | cycle length | 28.2 | 2.6 | 23 |  |  |  |  | 34 |
| 1992 | France |  |  | 139 | 1060 | ovulation | 15.5 | 3.1 | 9 |  |  |  |  | 22 |

### Privacy policies of the two applications

**Privacy Policy**

| Organisation | Sympto – Fondation Symptothem |
| --- | --- |
| General website | https://sympto.org/home-main_en.html |
| Link to the privacy policy | https://sympto.org/legal_en.html |
| Retrieved on | November 24^th^ 2017 |
| Version of document | 1.3 |
| Available languages | French, English, German, Italian, Spanish, Polish, Russian, Bulgarian, |

**General conditions**^[1]^

**sympto features:**

- The *sympto* app (TM) is not a medical device but an electronic learning tool that enables healthy women to take charge of their fertility in all gynecological situations: fertility awareness (ovulation), effective, ecological contraception, also during breastfeeding and pre-menopause, as well as pregnancy achievement and monitoring during pregnancy.
- Information on this site is not meant to substitute for the advice of a physician or medical professional and should not be not used for diagnosing or treating a health problem or disease, or prescribing any medication. Information and statements regarding dietary supplements have not been evaluated by the Food and Drug Administration and are not intended to diagnose, treat, cure, or prevent any disease
- The SymptoTherm Foundation offers counseling that will guide you on your path to body literacy. We do recommend you study the Manual 'The Complete Symptothermal Guide' avalable on our website for download free of charge. If you do not want to study the Learning Manual on your own, you have the option of contacting a symptothermal counselor. There will be a fee set by her for this service.
- If you choose unlimited access to sympto.org, there will be no consultation service included. 'Unlimited' means that the access on your sympto.org account is guaranteed as long as the SymptoTherm Foundation can uphold this service. In case of an accident within our principal server, we decline all liability. However, your data is regularly backed up on a second server. If you do not use your unlimited sympto.org account for more than one year, sympto.org is entitled to demand a reactivation fee before use or cancelation of your account. An unlimited account cannot be transferred: sympto.org will cancel all unlimited accounts whenever the email address has changed, unless this change has been approved by sympto.org. The client cannot delete her account because of possible abuse but she can ask sympto.org to deactivate it. The SymptoTherm Foundation does not sell any of your data to third parties.
- If you lose your cell phone, all data which you have transferred and thus saved to your sympto.org account can be retrieved. All your data is treated confidentially. Sync your data weekly to have it backed up in your web account. You may incur transmission costs for synchronization if you are using an older generation device. Theses costs are negligible and displayed in your account.
- Your sympto.org account provides you with a full chart of your cycle. The cell phone chart cannot display the complete representation of all your cycle data nor can it offer all the features of your home account on sympto.org.
- We recommend that you print out your charts periodically as soon as your cycle is completed. The charts provide you with a permanent record of your gynecological history. You can send them to whoever you like for discussion and analysis either as an attachment or via a internet link that you have to activate in the settings of your account. The chart is the only serious basis for discussion whenever you are concerned about irregular cycles or other problems. It contains valuable information.
- While keeping your identity totally confidential, the sympto.org website may use your data for statistical purposes in order to improve knowledge of the female cycle and to disseminate the effectiveness of the STM.

**Additional service features if you are pregnant or breastfeeding**

Whenever you suspend your observations because of a pregnancy or any other reason, your contract continus according to your payment. *sympto* is still useful as it can become your pregnancy diary. it also indicates the probable day of your delivery.

When your contract expires, your cycles start being deleted as follows: the system always stores the previous 360 days. If, for example, you resume your observations on the 11th month after a pause of several months, your last cycle will still be there and the data stored in you cycle chart will serve as a basis for the future. But once a year has elapsed, we cannot guarantee you the storage of the files.

**Disclaimer**

No woman or couple can ever be absolutely certain of avoiding an unwanted pregnancy. Even the best contraception can fail. In such a case, the woman or the couple has to decide on their own whether they want to take responsibility for the child. We do not accuse anybody of actively promoting abortion. Teenagers who do not have regular biphasic cycles can use sympto,ch to improve their fertility awareness (knowledge about ovulation) but not for contraceptive purposes.

sympto.ch does not protect you against sexually transmitted diseases. The sympto.ch system has been developed by Symptotherm Foundation, according to article 4 of its Foundation chart, in order to improve and facilitate the learning process and make the use of the STM more attractive. The sympto.ch system makes cycle observations easier but it does not make them redundant: the woman remains the only person responsible for her observations. F. i. she must know how and when she has to discard problematic temperatures. The SymptoTherm Basics booklet contains the elementary information about the female cycle needed to fully understand the User's Manual and the FAQ of this website.

Symptotherm Foundation can change the User's Manual (Learning Manual) at any time without any previous warning according to the system?s adaptations to its environment. 

The fertility indications processed by sympto.org all aim at the same goal: facilitating and enhancing women's self-observations. The sympto.org system offers an output of the information that the woman has previously entered. She fully takes charge of her input. She also can and sometimes has to change past values according to the STM principles and practice, but she endorses full responsibility for her modifications.

**Scientific records**

The symptothermal laws describe the biological processes of the female cycle and are explained our Manuals and in any symptothermal guide. The principle of cross-checking at the beginning and the end of the fertility span are based on the internationally recognized German and Austrian NFP/INER schools. The double system of internal and external cervical fluid observation is inspired by A Couple's Guide to Fertility, The Complete Symptothermal Method, Rose Fuller and Rev. J. Huneger, Northwest Family Services, Portland, Oregon. (This book is the American extension of the INER approach developed by J. Rötzer.) The option allowing the user to focus either on the temperature signs only or on the cervical fluid observations (Billings mode) is an innovation of our Foundation, as is the interchangeability of the vaginal sensation level and the cervix examination. We also have reformulated the symptothermal laws in our Manuals and introduced into *sympto* educational tools for beginners (motivation messages, daily recommendations, error messages). The entry board for the daily observations as well as the design of cycle charts are our copyright.

When used correctly, the *sympto* fertility indicator will not display 'false negative' results. This means that the system does not display 'infertile' when the cycle is still in its fertility window, according to the symptothermal standards. 'Correct use' means that the observation quality of the woman has been validated out of at least 6 cycles on sympto: either she has introduced 6 of her former cycles when opening an account (which are checked by the sympto administration) or by 6 new learning cycles, monitored by a sympto counsellor. Thus, the risk of an unwanted pregnancy is eliminated. *sympto* application is very close to the perfect 'classic' use, i.e. comparable to the competent and conscientious manual user. However, *sympto* displays false positive results in the beginning of the cycle just like the classic NFP use. This means that for the sake of effectiveness, it declares certain days as fertile which in fact are not. The number of false positive days will decrease after 12 cycles of observation, again according to the German NFP-standards. It is however possible that *sympto* might not be able to detect all temperature rise scenarios, especially if they are exceptional and complicated. These exceptions will generate more false positive days within the cycle.

**Terms and conditions for purchasing books and thermometers**

Your order is processed within the next 3 working days after your purchase transaction. The product is shipped by ordinary mail within the next 7 working days. The shipment destination is the address indicated with your order. In the event that the product is not received due to a postal error, SymptoTherm Foundation does not endorse any liability. There will be no refund if SymptoTherm can prove that the product has been sent correctly. Shipment and packing costs are indicated for each article. Not included in these costs are duty fees or local taxes.
Your payment has SSL security according the safety standards of Saferpay SA.

**Privacy of sympto.ch**

The sympto.ch website guarantees the perfectly confidential management of all your personal cycles. It presents your cycles on coloured and styled chart that can be printed out. It investigates the statistics in order to improve the application *sympto* and it encourages you to download the up dates. sympto.ch guarantees to keep the stock of at least your last 12 cycles or the last 360 days. Even if you do not synchronize during several months you can start all over again or continue as long as your account is active.

Without being asked, sympto.ch administrators avails himself of the right to check your cycles and to make any suggestions that might improve your cycle observations. The traceability of all modifications made by the user on her cycle chart is guaranteed. The counsellor can supervise these modifications in the back office. The service quality is thus fulfilled as well as the conditions for making scientific or clinic studies.

**Copyright 2008**

Unless otherwise indicated, all rights to this website and the information therein, including copyrights and other intellectual property rights, are owned by Symptotherm Foundation. The domain name sympto.*** is an intellectual property of Symptotherm Foundation and cannot be used as a site name which promotes natural conception regulation (natural family planning) or Fertility Awareness.

The formula which are integrated into the *sympto* soft ware are relatively easy and based on well known biological laws. However, the design of the sympto.ch, which also takes fully into account irregular and non ovulatory cycles, is very complex and protected by the copy right of Symptotherm Foundation. The copy right encloses the following elements: Fertility indication, the *sympto* engine and its algorithms, the sympto.ch site, the icon language on the cycle chart and on the overall site. This enumeration is not conclusive. 

The code source of sympto is available on request and for free as long as the mission of the Foundation SymptoTherm is respected.

There is now an open source version of the sympto code available on demand.

Users are permitted to read the website and the information therein and make copies for their personal use, for example by printing or storing it. All other uses of the website or the information, for example the storage or reproduction of (a part of) the website of Symptotherm Foundation in any external internet site or the creation of links, hypertext links or deeplinks between the website of Symptotherm Foundation and any other internet site is prohibited without the express written consent of Symptotherm Foundation. We are not liable for the contents of linked sites and pages.

**Privacy Policy**

| Organisation | Kindara |
| --- | --- |
| General website | https://www.kindara.com/ |
| Link to the privacy policy | https://www.kindara.com/privacy-policy |
| Retrieved on | November 24^th^ 2017 |
| Version of document | 1.2 |
| Available languages | English |

### Privacy Policy - KINDARA^[2]^

#### EFFECTIVE DATE: JUNE 21, 2011

At Kindara Inc. (“Kindara”), your privacy is important to us. This Privacy Policy (the “Policy”) provides you with information about how we may collect, use, and share your personal information so that you may make knowledgeable choices about the information that you choose to provide to us. “Personal Information” includes information that alone or when in combination with other information may be used to readily identify, contact, or locate you.

Please note that we may update this Privacy Policy from time to time, so please check back with us periodically.

This Privacy Policy applies to the Kindara services made available via applications on mobile or handheld devices (the “App”s), the Internet and our website and web application located at www.kindara.com (the “Website” and “Web App” respectively) (collectively, the “Service”). When you use the Service, you consent to Kindara’s collection and use of information from or about you as well as information related to your use of the Service.

This Privacy Policy provides information regarding Kindara’s data practices, including:

- How Kindara collects personal information
- How Kindara uses “cookies” and related information collected automatically when you use our Website or App
- How Kindara uses personal information
- What personal information, if any, Kindara may share about you
- Security measures that Kindara employs to protect your information from accidental loss or disclosure
- Other things you should know about privacy and Kindara

#### COLLECTED INFORMATION AND USAGE

Kindara collects and uses the information you provide to us when you use the Kindara Service. Information that Kindara may collect includes: name, date of birth, e-mail address, fertility-related data and other family planning and health-related information you provide. You may consider some of this information to be sensitive so you should choose carefully regarding whether and if you will use the Service.

We may use or share your Personal Information (e.g., name, address, telephone number, email address, and location) where it is necessary for us to complete a transaction or do something that you have asked us to do.

How Kindara collects and uses information from you:

- Website, Web App, or App Usage. Kindara (or its vendors and suppliers) may observe your activities, preferences, and transactional data (such as your IP address and browser type) as well as content you have viewed during your use of the Service. We may use this data for any purpose unless we tell you otherwise in connection with a particular Service. While we may collect or log this information, we do not identify you or match this non-Personal Information with your other Personal Information.
- Customer Service. If you purchase a paid Service offered by Kindara you may be asked for information needed to complete your request and provide related customer service. This information you provide is used, for example, to process your transactions, to create and share aggregate reports about these transactions, and for any risk management functions relating to transactions. If you purchase a subscription Service, your transaction may be conducted through our vendors and suppliers. Vendors and suppliers engaged by Conscious may have their use of your Personal Information limited by this Privacy Policy, contractual restrictions, and applicable law. Kindara (through its vendors or suppliers) may limit its provision of services to particular jurisdictions and as permitted by applicable law.
- Development of the Kindara Service; Scientific Research. In our mission to better understand fertility cycles and help women make powerful and informed choices around their fertility, Kindara will use your fertility data anonymously in our own research to better understand human fertility and improve our products and services. We may also share anonymous Personal Information with third-party researchers. Personal Information we share with a third party will be anonymous and not associated with your name or any identifying information provided by Kindara.
- Technical and Administrative Support. Kindara may allow you to provide registration information to us associated with a particular Kindara Service or products. Kindara may also use the information you provide in online registration forms to notify you periodically about important changes to the Service (such as a change to this Privacy Policy or notice of a security breach), and new products and services and related features. If you register, we may ask for, among other things, your name, address, email, telephone number, type of product purchased, or other product-identifying information.
- Promotions, and Contests. You may be asked to provide Personal Information, for entry into a particular promotion or contest, so that we can let you know if you won a prize. The specific rules and regulations governing the particular promotion or contest will vary and your participation constitutes your agreement to abide by those rules and regulations.
- Marketing. When you provide Kindara with Personal Information, we may communicate with you using the information you have given us to provide you with information we think may be of interest to you including marketing and promotional messages. For special offers and promotions we will provide you with an opportunity to opt-out of receiving further similar communications.

#### NON-PERSONAL INFORMATION, COOKIES, AND RELATED INFORMATION COLLECTION

Cookies are small lines of text/data that are written onto your computer by a website. The Service may involve use of cookies or other similar technologies to collect and store non-Personal Information. We utilize these technologies to research and understand how the Service is used, to develop and improve the Service, to personalize your online experience, and for advertising and other marketing purposes.

When you exit out of the Service, session cookies used by Kindara are removed . In some instances, such as where you register in order to customize your use of the Service, cookies may persist so that we can recognize you without requiring log-in at each visit. These persistent cookies placed in connection with your use of the Service may remain stored on your computer or mobile device until you remove them. Disabling or removing cookies may require you to manually log on each time you wish to use the Service.

#### WHEN KINDARA MAY SHARE COLLECTED INFORMATION

In addition to the circumstances noted above, Kindara may also share your Personal Information with third parties in the following situations:

- Enabling Services. Kindara contracts with other companies in order to enable provision of the Service. For example, web based services require web hosting, thus Kindara may contract with web hosting service providers.
- Public Areas. The Service may include forums for communicating with others; such forums are public areas. Any information disclosed in such public areas of the Service or on any other website (Facebook, YouTube, Twitter, etc.) will become public information. You should exercise caution when disclosing information in these public areas, especially your health-related and other sensitive information. Content posted in public areas of the Service, including advice and opinions, represent only the views and input of the poster; Kindara does not necessarily endorse, support, verify, or agree with such content. If you have any questions or comments about any content on the Service please send us an email at:
- Business Changes. If Kindara should ever file for bankruptcy or merge with another company, or if Kindara should decide to buy, sell, or reorganize some part or all of its business, Kindara may be required to disclose your Personal Information to prospective or actual purchasers in connection with one of these transactions.
- As Required by Law and Other Extraordinary Disclosures. Kindara may be required to disclose your Personal Information if it: (i) believes it is reasonably necessary to comply with legal process (such as a court order, subpoena, search warrant, etc.) or other legal requirements of any governmental authority, (ii) would potentially mitigate our liability in an actual or potential lawsuit, (iii) is otherwise necessary to protect our rights or property, or (iv) is necessary to protect the legal rights or property of others.

#### YOUR RIGHT TO ACCESS, MAKE CHANGES TO, OR REQUEST DELETION OF PERSONAL INFORMATION

You may request access to, correction of, or deletion of your Personal Information. For instructions on changing any of your privacy preferences, accessing your information, updating your information or for any privacy or data-protection-related question please write to for assistance.

#### SECURITY

Data Transmission. No data transmissions over the Internet can be guaranteed to be 100% secure. Thus, Kindara cannot ensure or warrant the security of any information that you transmit to us. You use the Service at your own risk. Once we receive your transmission, we make reasonable efforts to ensure security on our systems. However, this is not a guarantee that such information may not be accessed, disclosed, altered, or destroyed by breach of such firewalls and secure server software. In the event of a security breach. If Kindara learns of a security system’s breach, we may attempt to notify you electronically so that you can take appropriate protective steps. By using the Service or providing Personal Information to us, you agree that we can communicate with you electronically regarding security, privacy, and administrative issues relating to your use of the Service. Also, Kindara may post a notice on the Website or the App if a security breach occurs. If this happens, you will need a web browser or mobile device enabling you to view the Service. Kindara may also send an email to you at the email address you have provided to us in these circumstances. Depending on where you live, you may have a legal right to receive notice of a security breach in writing. To receive free written notice of a security breach (or to withdraw your consent from receiving electronic notice) you must notify us at.

#### OTHER IMPORTANT INFORMATION

- Children. Kindara Services are not directed at children. By using the Service, you represent and warrant that you are 18 years of age or older. If a person under 13 submits information through the Website or the App, and we learn the person submitting the information is a child, we will attempt to delete this information as soon as possible. Because we do not knowingly collect any Personal Information from persons under 13, we do not use or disclose such information to third parties.
- Third party websites. The Service may contain links to third party websites. Such third party websites are not governed by this Privacy Policy. If you choose to visit such third party websites, you should review the third party’s applicable privacy policy carefully.
- Data Processing in the United States. Kindara is located in the United States. The Service is hosted in the United States, and directed at residents of the United States and Canada only. By using the Service you consent to the processing of your Personal Information in the United States and elsewhere as described in this Privacy Policy.

#### CHANGES TO THIS PRIVACY POLICY BY KINDARA

Kindara reserves the right to change this Privacy Policy in the future. If we make changes to this Privacy Policy that are materially less restrictive on our use of personal information than we have previously collected we will make reasonable efforts to obtain your consent to provide notice and obtain your consent to any such uses. We will post the updated Privacy Policy through the Service. Your continued use of the Service following a change in the Privacy Policy represents your consent to the new Privacy Policy to the fullest extent permitted by law. We encourage you to periodically review this Privacy Policy.

#### CONTACTING KINDARA

Kindara Inc. Box 3801630 30th St. Unit A Boulder, CO 80301

1. Retrieved on 24.11.2017 from https://sympto.org/legal_en.html
2. Retrieved from https://www.kindara.com/privacy-policy
